# Conformational Remodeling Underlies Activity Loss in Disease-Linked Asparagine Synthetase Variant

**DOI:** 10.64898/2026.02.01.703106

**Authors:** Lucciano A. Pearce, Adriana Coricello, Alanya J. Nardone, Giovanni Bottegoni, Yuichiro Takagi, Wen Zhu

**Affiliations:** Department of Chemistry and Biochemistry, Florida State University, Tallahassee, FL, U.S.; Department of Biomolecular Sciences, Università degli Studi di Urbino Carlo Bo, Urbino, Italy; Department of Pharmacy, University of Birmingham, Birmingham, U.K.; Department of Biochemistry, Molecular Biology and Pharmacology, Indiana University School of Medicine, Indianapolis, IN, U.S.

**Keywords:** asparagine synthetase deficiency, rare disease, pathogenic mutation, enzyme kinetics, cryogenic electron microscopy, molecular dynamics, conformational dynamics

## Abstract

Asparagine synthetase deficiency (ASNSD) is a devastating congenital disorder characterized by profound neurological impairment and early childhood mortality. It is associated with pathogenic mutations in the asparagine synthetase (ASNS) gene. Despite the critical role of ASNS in the amino acid cycle, the molecular basis by which ASNSD-linked missense mutations impair enzyme function remains poorly understood. Here, we present a comprehensive characterization of a recurrent ASNSD-linked variant, R48Q. Steady-state kinetic assays reveal severe reductions in L-glutamine-dependent catalysis and disrupted product stoichiometry, implicating impaired interdomain communication. Cryogenic electron microscopy (cryo-EM) and 3D variable analysis of the EM map uncovers altered loop conformations at the N-terminal active site and subtle conformational changes at the C-terminal domain. Consistent with the structural data, molecular dynamics simulations support that the local disruption propagates across the protein, thereby decoupling coordinated domain motions essential for catalysis. Additionally, we demonstrate that the flanking arginine and the affected loop are evolutionarily conserved across Class II glutamine amidotransferases, highlighting their shared mechanistic importance. These findings provide the molecular basis of an ASNSD variant and establish a framework for understanding how point mutations disrupt complex enzyme dynamics, with broad implications for precision medicine.

**Significance:** Understanding how mutations affect multidomain enzymes is crucial for elucidating the molecular mechanisms underlying genetic disorders. Here, we examine the molecular consequences of the R48Q variant in human asparagine synthetase (ASNS), the sole enzyme responsible for *de novo* L-asparagine synthesis; mutations of this enzyme lead to a fatal neurometabolic disorder, asparagine synthetase deficiency (ASNSD). By combining biochemical, cryogenic electron microscopy, and molecular dynamics simulation, we show that a single N-terminal amino acid substitution disrupts both local and global coordination, impairing enzyme activity. Our work provides the first mechanistic blueprint of an ASNSD-linked variant. These findings not only deepen our understanding of ASNS but also offer a generalized framework for studying the dynamic regulation of multidomain enzymes in disease.

## Introduction

Nearly 1500 human enzymes are associated with over 2500 clinically identified inherited diseases, most of which are rare diseases and have no effective treatment due to a limited understanding of the molecular mechanism (1, 2). A large fraction of these diseases arises from missense mutations, in which a single nucleotide change results in an amino acid substitution throughout the enzyme’s structure (3–5). In the absence of experimental validation, bioinformatic annotations oversimplify the impact of enzyme variants, attributing changes in activity mostly to altered thermostability without further mechanistic explanation (6). This generalization has hindered efforts to understand the molecular mechanisms underlying how and why these residue substitutions impair biological function, thereby hindering the development of treatments targeting the root cause of these diseases.

Asparagine synthetase deficiency (ASNSD) is an inborn neurometabolic disorder (7, 8). Individuals affected by ASNSD exhibit severe neurological impairments, including cognitive delays, axial hypotonia, intellectual disability, progressive cerebral atrophy, and intractable seizures (7–10). Whole-genome sequencing of individuals affected by ASNSD has linked ASNSD to homozygous or compound-heterozygous mutations in the gene encoding asparagine synthetase (ASNS) (11–13). Diagnosis is often supported by abnormal neuroimaging and reduced L-asparagine levels in cerebrospinal fluid and/or plasma, although these metabolites are not consistently low in all affected individuals (9, 14–16). As a result, L-asparagine supplementation has been explored as a potential strategy to improve or reverse symptoms (7, 14). The outcomes of L-asparagine supplementation, however, vary considerably. Some reports indicate only mild developmental improvements, including increased attention and enhanced nonverbal communication, whereas others describe a worsening of symptoms, highlighting the poorly understood and potentially mutation-specific molecular mechanisms underlying ASNSD. (14, 17).

Most ASNSD-linked ASNS variants are single-residue substitutions throughout the protein (**SI Appendix, Table S1**) (18). Among the ASNSD-linked variants, the R48Q (NM_001673.5:c.146G>A) substitution has been recurrently identified in multiple individuals with severe neurological symptoms (15, 19). Their fibroblasts showed impaired growth in L-asparagine-depleted media, suggesting damaged asparagine production in the cells (15). Since the ASNS expression level and structural integrity remain unaffected, the substitution is proposed to affect enzymatic activity by impairing L-glutamine binding, as the mutation is located in the N-terminal domain (15, 20). However, the molecular mechanism by which the R48Q substitution impairs asparagine production has not been experimentally tested.

Asparagine synthetase (ASNS) catalyzes the ATP-dependent synthesis of L-asparagine through two spatially separated catalytic domains (**Fig. 1a**) (18, 21). In the N-terminal glutaminase domain, L-glutamine is hydrolyzed to L-glutamate and ammonia, a reaction initiated by the conserved N-terminal residue Cys1, which serves as the catalytic nucleophile (**Fig. 1a**) (22). Structural insight into the role of Cys1 has been provided by an ASNS structure covalently modified by the glutamine analog 6-diazo-5-oxo-L-norleucine (DON), which traps the enzyme in a catalytic intermediate state prior to glutamine hydrolysis (**Fig. 1b**) (23). In this DON-modified structure, Cys1 forms a covalent adduct with the inhibitor, and a loop spanning residues Arg48– Met58 (Loop 1) adopts a closed conformation that positions key residues in close proximity to both Cys1 and the bound intermediate (**Fig. 1b**). These observations suggest that conformational coupling between Loop 1 and Cys1 plays a critical role in organizing the glutaminase active site for efficient catalysis. Once ammonia is generated, it is then channeled through an ∼20 Å intramolecular tunnel to the C-terminal synthetase domain, which participates in the amidation step (24–26). In the C-terminal domain, ATP is coordinated by magnesium ions and activates the carboxylate group of L-aspartate, generating inorganic pyrophosphate and a β-aspartyl-AMP intermediate (27–29). The translocated ammonia subsequently attacks this intermediate via nucleophilic substitution, yielding AMP and L-asparagine as the final products (23, 30).

**Figure 1.**
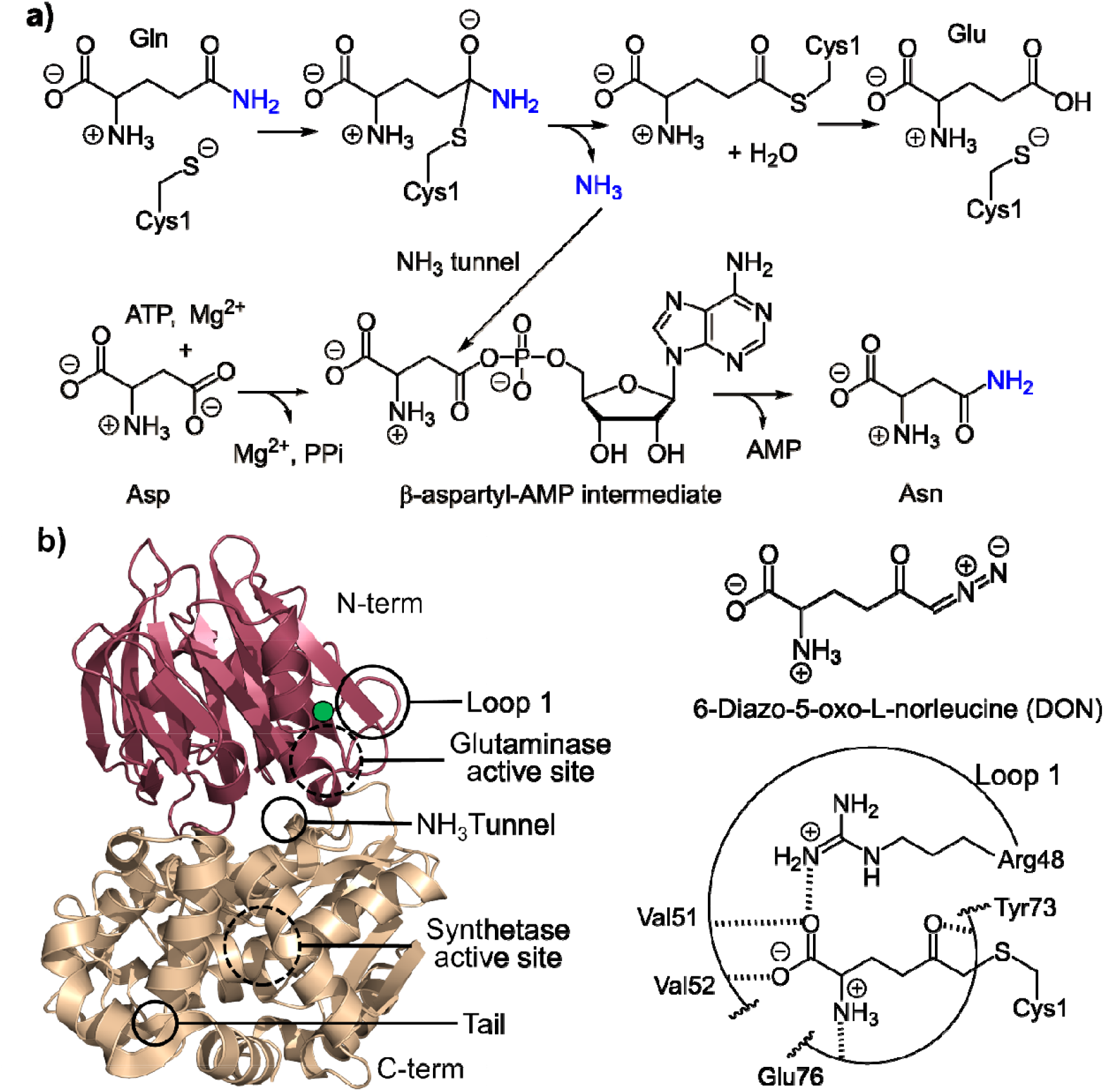
Bifunctional mechanism of human asparagine synthetase (ASNS). a) ASNS catalyzes the ATP-dependent biosynthesis of L-asparagine from L-aspartate and L-glutamine through a two-domain, two-step reaction. Top raspberry box: In the N-terminal glutaminase domain, L-glutamine (Gln) is hydrolyzed to L-glutamate (Glu) and ammonia (NH_3_), which is then channeled through an ∼20 Å intramolecular tunnel. Bottom tan box: In the C-terminal synthetase domain, ATP and Mg^2+^activate L-aspartate (Asp) to form a β-aspartyl-AMP intermediate, which reacts with the translocated NH_3_ to generate L-asparagine (Asn) and AMP. b) The crystal structure of the human ASNS is shown on the left: the N-terminal glutaminase domain in raspberry and the C-terminal synthetase domain in tan. Black dashed circles indicate the two active sites, and the conserved residue R48 is marked with green dot. The chemical structure of DON and the polar interactions (dashed line) identified in the N-terminal active site of DON-modified WT are shown on the right.

In this study, we investigated the molecular consequences of the R48Q substitution through integrated enzyme kinetics, cryogenic electron microscopy (cryo-EM) and molecular dynamics (MD) simulations. We uncover how this single residue change impairs enzyme structure, function, and dynamics at both local and global levels. Our findings reveal that Arg48 is essential for maintaining the N-terminal active-site loop in a catalytically competent conformation, thereby enabling productive engagement of L-glutamine. We also identified that the conformational perturbation introduced by the R48Q variant propagates beyond the glutaminase site, ultimately affecting the C-terminal domain of ASNS. This work provides the first atomic-level mechanistic explanation for an ASNSD-associated variant and establishes a generalizable framework for understanding pathogenic missense mutations in other multidomain human enzymes.

## Results

### L-Asparagine production is impaired in the ASNS R48Q variant

#### The R48Q variant retains thermostability under physiological conditions

We prepared recombinant human ASNS WT and the R48Q variant using the baculovirus expression system, following our established protocol (23, 30). Consistent with the previous prediction (15, 20), the stability of the R48Q variant is only marginally impaired compared to that of the WT, as indicated by its unchanged circular dichroism profile at the physiologically relevant temperature and a marginal decrease (2.5 ^°^C) in melting temperature. (**SI Appendix, Fig. S1**)

#### Significant reduction in pyrophosphate production in the R48Q variant compared to WT when L-glutamine serves as the nitrogen source

To determine the steady-state kinetics of the R48Q variant, we adapted two kinetic assays to independently monitor the production of inorganic pyrophosphate, L-glutamate, and L-asparagine (23, 24, 31). Under physiological conditions, L-glutamine is the biologically relevant nitrogen source, while the C-terminal synthetase active site of ASNS can directly utilize ammonia from solution in the *in vitro* assays (25). Therefore, kinetic measurements were conducted using either L-glutamine or ammonia as the nitrogen source to evaluate the impact of N-terminal residue substitution on the function of each domain individually. Our kinetic assays reveal the *k*_*cat*_ for pyrophosphate production decreased approximately 50-fold in the R48Q variant using the UV-Vis-based real-time kinetic assay (**Table 1** and **SI Appendix, Fig. S2**). While the apparent K_M_ (^app^K_M_) for L-glutamine in the R48Q-catalyzed reaction remains comparable to that of WT, the ^app^K_M_ values for ATP and L-aspartate decreased by 8- and 10-fold, respectively. Consequently, the catalytic efficiency (*k*_*cat*_/^app^K_M_) is reduced by 37-, 5-, and 17-fold for L-glutamine, ATP, and L-aspartate, respectively. The most pronounced *k*_*cat*_/^app^K_M_ impact is on L-glutamine, suggesting that N-terminal glutaminase activity is severely compromised in the R48Q variant.

**Table 1.**
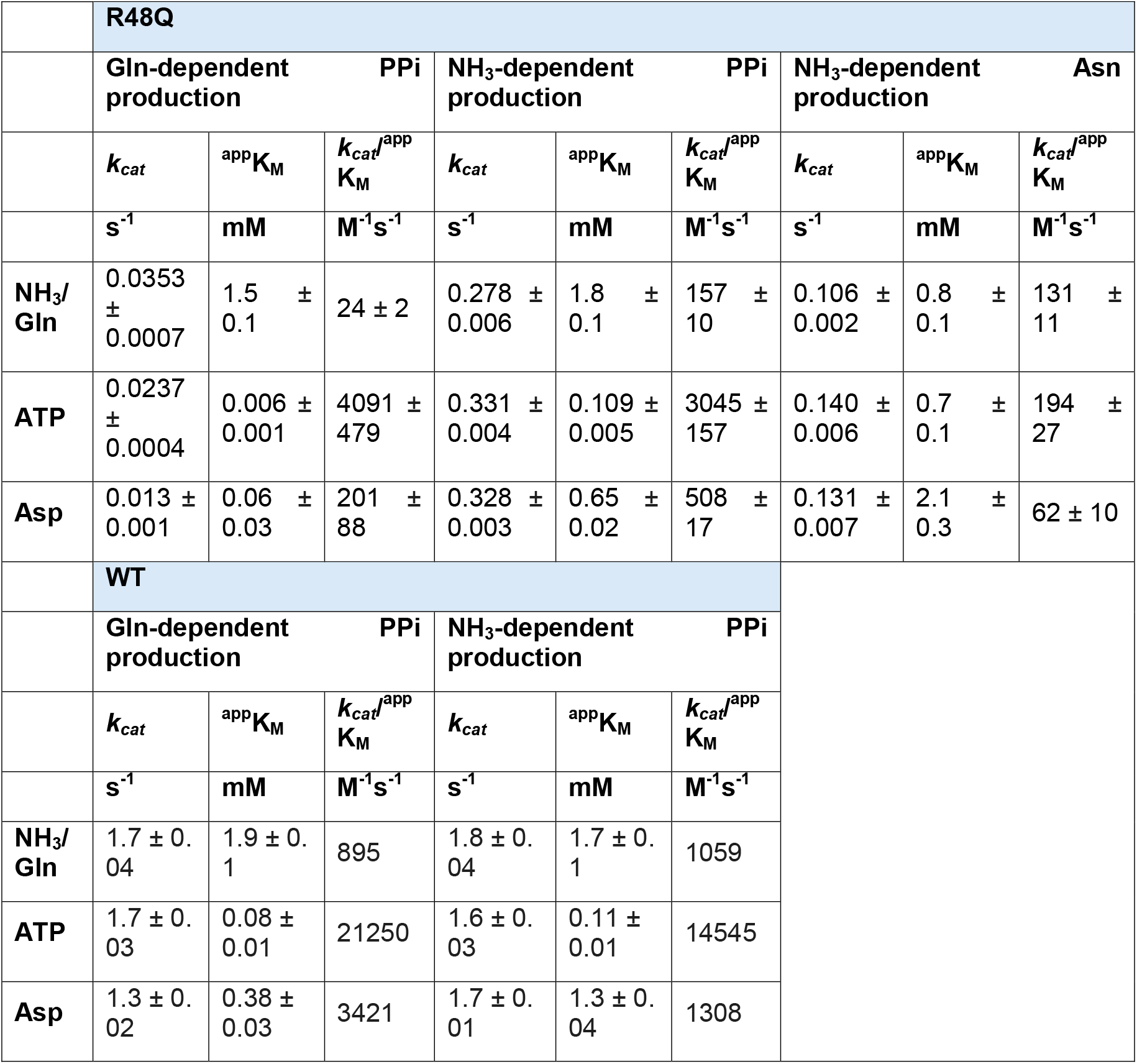
Steady-state kinetics of the ASNS R48Q variant. Kinetic parameters for each substrate were determined for the ASNS R48Q variant using either L-glutamine (Gln) or ammonia (NH_3_) as the nitrogen source. Pyrophosphate (PPi) production was measured using a UV-vis-based enzyme-coupled continuous assay, and L-asparagine (Asp) production was assessed using an HPLC-based stopped assay. For comparison, WT ASNS values obtained from the reference (78) are shown at the bottom. All values are reported as the means of independent duplicates ± standard error.

#### Activity in the C-terminal synthetase domain is also impaired in the R48Q variant

ASNS is a bi-domain enzyme, composed of a glutamine-hydrolyzing domain and a synthetase domain (23, 29). It is known that the activity of the two active sites can influence each other in many other bi-functional enzymes of this type (32–35). To further distinguish the impact directly caused by the dysfunctional N-terminal active site from the potential secondary impact that occurs in the C-terminal active site or the ammonia trafficking, we assessed pyrophosphate production when ammonia was directly supplied in bulk solution (**Table 1** and **SI Appendix, Fig. S2**). Under these conditions, although the ^app^K_M_ values for ammonia, ATP, and L-aspartate remained unchanged, the *k*_*cat*_ of the R48Q variant is one-fifth that of WT ASNS. These findings indicate that substrate engagement at the C-terminal active sites is also altered by the Gln48 substitution, although to a lesser extent than the disruption at the N-terminal active site. Furthermore, the identical ^app^K_M_ values for ammonia and glutamine-dependent activity in both WT and R48Q suggest that the fundamental catalytic mechanism of the glutaminase active site remains intact. Instead, the substitution primarily affects the concentration of catalytically competent enzymes during glutamine hydrolysis.

Next, we monitored L-asparagine formation under the same experimental conditions that were used to track pyrophosphate production (**Table 1** and **SI Appendix, Fig. S3**). The observed *k*_*cat*_ for all substrates is approximately 0.1 s^-1^, which is 3-fold lower than the values obtained from pyrophosphate production assays. Additionally, the ^app^K_M_ for ammonia decreases by 2-fold, while the ^app^K_M_ values for ATP and L-aspartate increase by 5- and 3-fold, respectively. This leads to 15- and 8-fold decreases in *k*_*cat*_/^app^K_M_ for ATP and L-aspartate, respectively. This data suggests that C-terminal synthetase activity is impaired by the removal of arginine in the R48Q variant.

Given that R48Q is located in the N-terminal domain, we next evaluated whether the residual glutaminase activity in R48Q can be modulated by substrate binding and catalysis at the C-terminal active site. We determined the L-glutamate and L-asparagine production when L-glutamine was used as the nitrogen source, both in the presence and absence of C-terminal substrates, ATP and L-aspartate (**SI Appendix, Fig. S4**). Our results show that while L-glutamate production in the R48Q variant is significantly lower than in WT, glutaminase activity is not influenced by whether catalysis occurs in the C-terminal active site.

We then calculated the pyrophosphate-to-asparagine production ratio across various substrate concentrations. In WT ASNS, pyrophosphate and L-asparagine are typically produced in a 1:1 ratio when either ammonia or L-glutamine serves as the nitrogen source (24, 36). Kinetic analysis of our *in vitro* assays, however, reveals a striking deviation in the R48Q variant, where the pyrophosphate-to-asparagine ratio is highly dependent on substrate concentrations (**SI Appendix, Fig. S3**). When ATP and L-aspartate are at saturating levels and ammonia concentration is varied, increasing ammonia raises the pyrophosphate-to-asparagine ratio from 1 to 3. When ammonia is saturated, lower ATP and L-aspartate concentrations favor an enzyme population that produces more pyrophosphate than L-asparagine, resulting in pyrophosphate-to-asparagine ratios of 15 and 10, respectively. As ATP and L-aspartate concentrations increase, the pyrophosphate-to-asparagine ratio decreases to 3. Under fully saturated substrate conditions, every three pyrophosphate molecules produced correspond to one productive L-asparagine formation when ammonia is used as the nitrogen source. When comparing L-glutamate and L-asparagine production during glutamine-dependent activity, we observed a 1:1 ratio in the R48Q variants, suggesting that the ammonia generated in the glutaminase active site can still be effectively used for L-asparagine production, similar to WT (**SI Appendix, Fig. S3**). Therefore, the ammonia tunnel is likely intact in the R48Q variant.

In essence, indicated by the series of kinetic analyses, the N-terminal active site of the R48Q variant is compromised in its ability to hydrolyze L-glutamine and generate ammonia. Moreover, the decoupling of pyrophosphate production from L-asparagine formation observed in the ammonia-dependent conditions suggests that the Gln48 substitution also disrupts the coordinated catalytic steps carried out in the C-terminal synthetase domain.

### Cryo-EM reveals specific structural perturbations in the R48Q Variant

To elucidate the structural basis underlying the kinetic effects observed in enzymatic activity assays, we determined the structure of the ASNS R48Q variant by single-particle cryo-EM. To enable high-resolution comparison between WT and the R48Q variant, we have collected cryo-EM datasets for both samples on a 300-kV Titan Krios operating in super-resolution mode. Data were processed using cryoSPARC v4.7.0, following a previously used strategy with modifications (see **Materials and Methods**) (23, 37). As outlined in the full processing workflows (**SI Appendix, Fig. S5 & S8**), particles were selected using template-based picking, followed by iterative rounds of 2D classification, *ab initio* reconstruction, and heterogeneous refinement to enrich for high-quality particle populations. Final reconstructions were obtained by non-uniform refinement, resulting in maps at 2.78 Å resolution for apo-WT and 2.81 Å resolution for the R48Q variant (**SI Appendix, Table S2** and **SI Appendix, Fig. S5-10**). Despite the improved resolution of the apo-WT ASNS structure to 2.78 Å, compared with the previously reported 3.5 Å model (30), several regions remain poorly ordered. In the final model, density is absent for two loop segments (residues 201-220 and 465-475) as well as for most of the C-terminal tail (residues 542-560), although two additional residues (W540 and I541) are now resolved. The lack of density in these regions suggests that they are disordered. A similar pattern of disorder is observed in the apo structure of the R48Q variant. Interestingly, Loop 1 (Arg48-Met58) is well resolved in the apo-R48Q structure. In contrast, in the apo-WT structure, EM density for several residues within this loop (Val51-Pro54) is weak, indicating increased flexibility in this region (**SI Appendix, Fig. S11**).

#### R48Q does not change the overall fold of ASNS

These high-resolution cryo-EM structures show that the R48Q variant maintains the overall fold of WT ASNS (**SI Appendix, Fig. S11**). The R48Q variant forms a head-to-head dimer with secondary structure features similar to those of the apo-WT (RMSD = 0.392 Å) structure and the DON-modified WT crystal structures (RMSD = 0.596 Å) (**SI Appendix, Fig. S12**). In both WT and R48Q, the dimer interface is stabilized by hydrogen bonds between anti-parallel β-sheets of the N-terminal domains (**SI Appendix, Fig. S12**). The absence of the intermonomer disulfide bond, as observed in the X-ray structure, confirms it is a crystallographic artifact (**SI Appendix, Fig. S13**). The two monomers in both WT and R48Q are nearly identical (RMSD = 0.212 Å for R48Q and 0.158 Å for WT). Key catalytic residues, such as Cys1 for glutamine hydrolysis in R48Q, remain in the catalytically active position within the N-terminal active site, as in apo-WT (**Fig. 2a**). Previous work from our group has shown that ASNS’s function depends on a tunnel-lining residue, Arg142, which acts as an essential gate for ammonia transport between the N-terminal and C-terminal active sites (30). In the R48Q variant, R142 maintains its closed orientation, as observed in the WT structure, consistent with kinetic data indicating that ammonia transport is not impaired (**SI Appendix, Fig. S13**).

**Figure 2.**
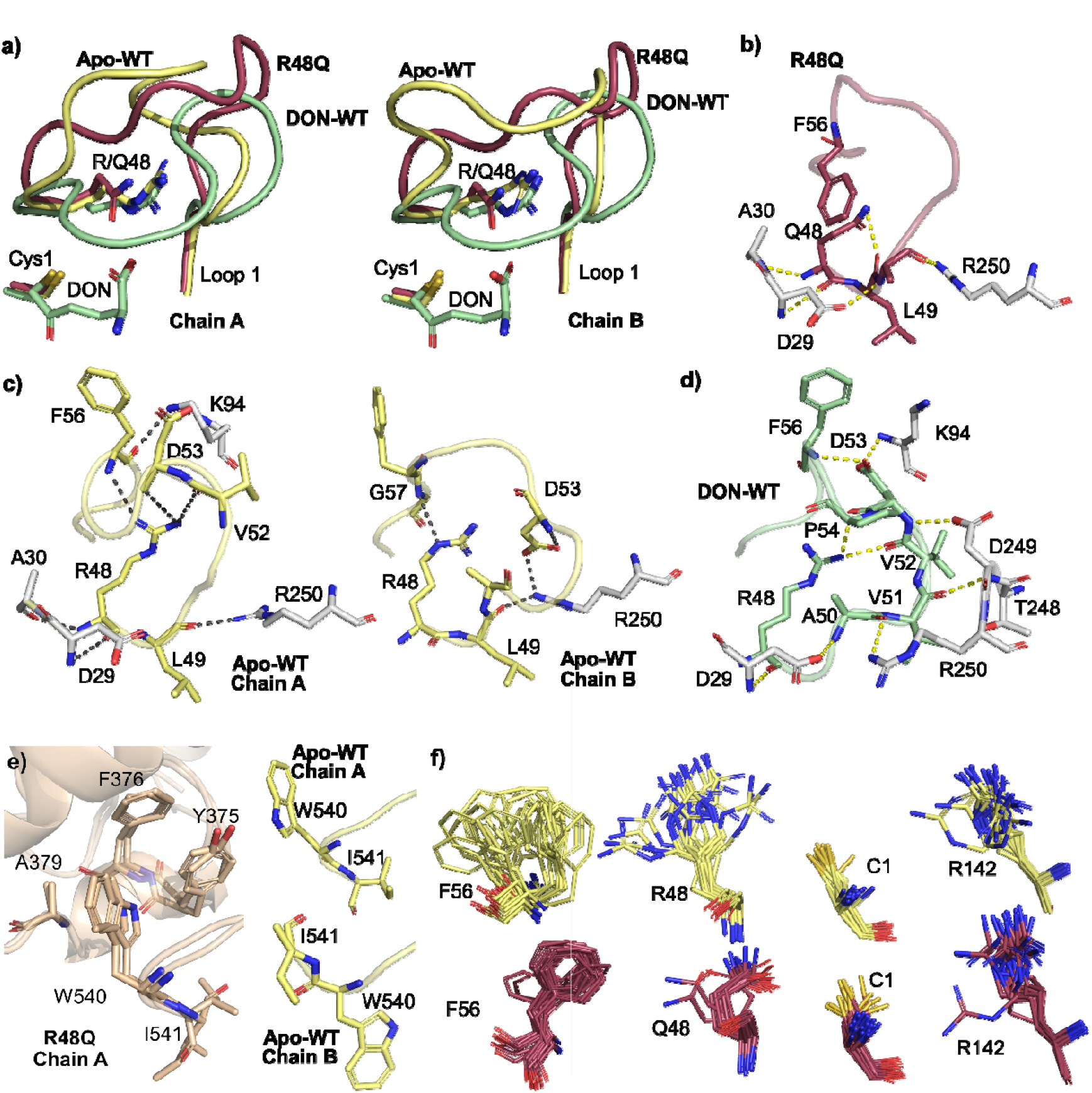
Cryo-EM structures of WT and R48Q ASNS reveal altered Loop 1 and C-terminal tail dynamics. a) Superposition of Loop 1 from the two monomers in the apo-WT (yellow), DON-bound WT (green), and R48Q (garnet) dimers, highlighting differences in backbone flexibility and side-chain positioning. b) Hydrogen-bonding networks surrounding Loop 1 in the R48Q variant. Only Chain A is shown, as the two monomers are structurally similar. The color scheme matches panel a; non-Loop 1 residues are shown in gray. c) Hydrogen-bonding networks surrounding Loop 1 in apo-WT, showing both Chain A and Chain B. The color scheme matches panel a; non-Loop 1 residues are shown in gray. d) Hydrogen-bonding networks surrounding Loop 1 in DON-bound WT. Only Chain A is shown, as the two monomers are structurally similar. The color scheme matches panel a; non-Loop 1 residues are shown in gray. Conformations of Trp540 and Ile541 in R48Q (wheat, left) and WT (yellow, right), demonstrating increased conformational heterogeneity in WT and restricted positioning in R48Q. 3DVA illustrating conformational heterogeneity of Phe56, Arg48, Cys1, and Arg142, colored as in panels a) and d).

#### R48Q alters the conformation of the N-terminal active site loop

As indicated before, conformational coupling between Loop 1 and Cys1 plays a critical role in glutaminase activity. Structural comparison of the R48Q variant, apo-WT, and DON-modified WT ASNS reveals that disruption of glutaminase activity in R48Q arises from altered conformational control of the catalytic residue Cys1 mediated by Loop 1. In the R48Q variant, Loop 1 consistently takes on an ‘open’ conformation in both monomers (**Fig. 2a&b**), while apo-WT shows significant flexibility in Loop 1 between its two monomers (**Fig. 2a&c**). The distance between Arg48 and the Val52 carbonyl fluctuates from 3.8 Å (chain A) to 7.1 Å (chain B), and the Arg48 side chain adopts different orientations in each monomer, suggesting the dynamic nature of Loop 1 in the WT enzyme. By contrast, in the WT ASNS crystal structure, where the catalytic Cys1 is modified by DON to mimic the covalent intermediate during glutamine hydrolysis, Loop 1 adopts a ‘closed’ conformation. This closed state is further stabilized by an extensive hydrogen-bonding network involving loop residues (Ala50, Val51, Val52, Asp53, Pro54 and Phe54), other N-terminal residues (Asp29 and Lys94), and C-terminal residues (Thr248, Asp249, Arg250), thereby structurally bridging the N- and C-terminal domains (**Fig. 2d**). In the DON-modified WT, Arg48 plays a central role in organizing the Loop 1 conformation via multiple hydrogen bonding interactions (**Fig. 2d**), it also forms a salt bridge to the DON carboxylate, thereby stabilizing the catalytically competent intermediate during catalysis (**SI Appendix, Fig. S14**). These observations indicate that Arg48 not only modulates loop conformation in apo-WT but also participates in stabilizing covalent intermediates during the catalysis. Therefore, substitution of Gln48 likely disrupts Loop 1 dynamics, impairing substrate entry and intermediate stabilization, thereby contributing to the observed loss of glutamine-dependent activity.

#### Subtle structural differences in the C-terminal active site of the R48Q variant

We compared the C-terminal active site of the R48Q variant and that of the WT ASNS, only minor changes in side-chain conformations were observed (**SI Appendix, Fig. S14**). For example, the side chain of Asp261 within the highly conserved SGGLDSS (PP-loop) motif, shared among many pyrophosphatases and involved in pyrophosphate binding, adopts distinct rotamer states in apo-WT, DON-bound WT, and R48Q. In DON-modified WT, Asp261 forms a salt bridge with Lys444 in both monomers (**SI Appendix, Fig. S14**). In the apo-WT structure, this salt bridge is absent due to the rotation of the Asp261 carboxylate. In R48Q, Asp261 samples multiple rotamers that resemble those observed in either the DON-modified WT or the apo WT. Both Asp261 and Lys444 are strictly conserved in ASNS across species, suggesting their functional importance (**SI Appendix, Fig. S15**). Similar rotamer variability is observed for Ser257 within the PP-loop when comparing the three structures. These conformational preference changes could affect ATP engagement and explain the lower apparent K_M_ for ATP and substantial reduction in *k*_*cat*_ in R48Q.

#### The WT C-terminal tail is more dynamic than that of the R48Q variant

Comparison of the two monomers in the WT and R48Q structures reveals a notable difference in the conformational behavior of Trp540 and Ile541. Trp540 and Ile541 are part of the functionally essential C-terminal tail of ASNS (16, 38), and Trp540 is highly conserved across ASNS homologs (**SI Appendix, Fig. S15**). In the WT enzyme, these residues adopt two distinct conformations, whereas in the R48Q variant, Trp540 and Ile541 are locked in the same position in both monomers (**Fig. 2e**). In the R48Q variant and in Chain A of the WT structure, Trp540 interacts with a hydrophobic pocket formed by Tyr375, Phe376, and Ala379 on a C-terminal domain helix. In contrast, in Chain B of the WT structure, the EM map clearly shows outward flipping of Trp540, accompanied by repositioning of Ile541 toward the hydrophobic patch (**Fig. 2e**). This conformational heterogeneity is consistent with the dynamic motion of Loop 1 observed in apo-WT and supports a model of coordinated structural remodeling.

### Three-dimensional variability analysis (3DVA) identifies an altered conformational ensemble in the R48Q variant

Since cryo-EM images contain conformational ensemble information of the enzyme (30, 39, 40), we performed 3DVA (41) on the R48Q variant, following our previously described methods for exploring the dynamic side chain rotations (30). We analyzed the EM map of R48Q and WT along five principal component axes, generating 100 high-resolution structures for each protein to identify the conformational ensemble present in the apo enzymes. The first and last frames in each cluster were presented as they correspond to the most structurally distinct conformations within each cluster (**Fig. 2f**).

#### Rotamer ensembles at the N-terminal active site shift in the R48Q variant

Our analysis revealed increased variability in the orientation of the catalytically essential Cys1 side chain in the R48Q structures compared to WT. In the WT ensemble, Phe56 and Arg48 sample a broad distribution of conformers, while the rotation of Cys1 remains relatively restricted. It’s been proposed that Cys1 must be deprotonated for catalysis (42, 43), this constrained orientation likely reflects the electrostatic influence posed by Arg48, helping to maintain a productive geometry for nucleophilic attack. In contrast, the R48Q variant exhibits reduced rotamer diversity at Phe56 and Gln48, while Cys1 samples a broader range of orientations than in the WT enzyme. The increased disorder in Cys1 highlights that the coordinated conformational coupling normally mediated by Arg48 in fine-tuning the geometry of the N-terminal active site is missing in the R48Q variant. The deviation of Cys1 in the R48Q from the catalytically competent geometry suggests a disruption of active-site preorganization, a feature essential for efficient nucleophilic attack during glutamine hydrolysis (32, 44–46). This conformational defect is consistent with the reduced catalytic efficiency observed in our glutamine-dependent kinetic analysis.

#### R48Q doesn’t alter the motion of ammonia tunnel-lining residues

In ASNS, we have previously identified Arg142 as a key residue that regulates ammonia trafficking (30). 3DVA analysis of the higher-resolution WT and R48Q structures demonstrates that Arg142 continues to adopt both open and closed conformations across both ensembles to a similar extent (**Fig. 2f & SI Appendix, Fig. S16 & S17**). Superimposition of the individual rotamers of other previously defined tunnel-lining residues, including Val119, Val141, Leu255, Met344, Val401, and Leu415, showed that these residues also adopt highly similar conformational assemblies in WT and R48Q (**SI Appendix, Fig. S18**). This conformational ensemble of tunnel-lining residues is fully consistent with our kinetic data, which indicates that ammonia trafficking remains unchanged in the R48Q variant.

### MD simulation captured altered global conformational dynamics

Observation of an alternative loop conformation in the cryo-EM structure of the R48Q variant prompted us to use MD simulation to examine changes in local and global dynamics in R48Q. We built a full-length WT ASNS model using ROBETTA and mutated residue Arg48 to glutamine (30, 47). The structure was equilibrated in a water box of the same size as used in the WT ASNS simulations (30). Four independent 200-ns simulations at 298.15 K were performed and analyzed.

#### Loop 1 exhibits altered dynamics in the R48Q variant

We first compared the dynamic fluctuations across the protein between WT and R48Q using RMSF values derived from the MD trajectories (**Fig. 3a & SI Appendix, Fig. S19**). The two proteins exhibit broadly similar fluctuation patterns, with only a few regions showing notable differences. The first region of interest comprises residues 49-52, which are part of Loop 1. Simulations reveal that the RMSF of these residues is higher for the R48Q variant than for WT, (**Fig. 3b & SI Appendix, Fig. S19**). The motion differences of Loop 1 can also be visualized in the distances between Asp53 and Lys94 and between Gln/Arg48 and Cys1, both of which are identified in the catalytically functional conformation of Loop 1 in the DON-WT structure (**SI Appendix, Fig. S20**). While the distance changes reflect a shift in conformational distribution in the first half of Loop 1 (residue 49-53), the magnitude of RMSF fluctuations in the second half of Loop 1 (53-58) remains similar between WT and R48Q, both around ∼2 Å. (**Fig. 3a & SI Appendix, Fig. S19**), thereby supporting the hypothesis that R48Q alters the ensemble of loop motion and partially stabilizes Loop 1 in a catalytically incompetent form.

**Figure 3.**
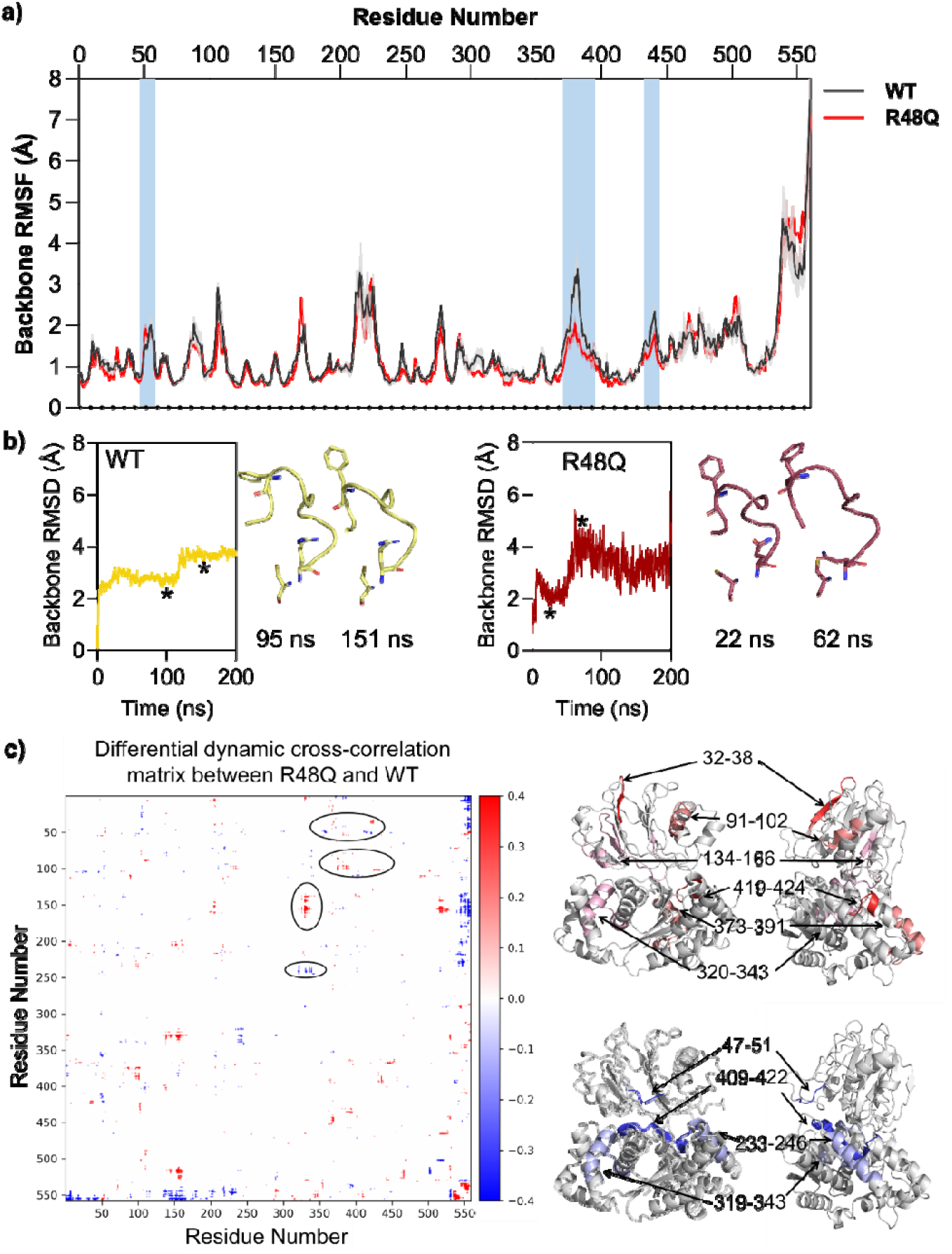
Fluctuation profiles from MD simulations reveal altered dynamics in the R48Q variant. a) RMSF traces for the backbone heavy atoms of WT ASNS (black) and the R48Q variant (red) highlight regions exhibiting altered flexibility in the variant. b) Backbone RMSD traces of the MD simulations of the Loop 1 region in WT (yellow) and the R48Q variant (garnet), along with the representative structures identified in the MD simulation corresponding to the asterisks in the trajectories. c) Difference between R48Q and WT hASNS dynamic cross-correlation matrices (DCCM). Only differences with an absolute value greater than 0.3 are displayed. In a difference DCCM (ΔDCCM), the color scale represents changes in correlated motion between two systems. Positive values (red) indicate an increase in correlated motion in the mutant compared to the reference. Negative values (blue) reflect a loss of correlated motion. Regions with values near zero indicate little to no change in dynamic coupling. The corresponding segments are mapped onto the R48Q structure on the right, with increased correlated motion shown in red and reduced correlated motion shown in blue.

#### Backbone fluctuation supports distal conformational change identified by cryo-EM

Interestingly, two regions in the C-terminal domain, residues 375-385 and 435-445, exhibited distinct fluctuation patterns between WT and R48Q (**Fig. 3a**). In both segments, the R48Q variant shows reduced fluctuations relative to WT. Structural mapping places these regions ∼50 Å from residue 48. Residues 375-385 contribute to the hydrophobic pocket that engages Trp540, and the dampened fluctuations in 375-385 align with cryo-EM observation that Trp540 is less dynamic in the R48Q structure (**Fig. 2e**). Residues 375-385, comprising a flexible loop and a short helix, packs directly against residues 375-385 through polar (Glu443-His378) and hydrophobic interactions (Ile437/Pro438-Ala379/Ala386) in the R48Q cryo-EM structure. Reduced fluctuations in both regions of R48Q suggest regional rigidification of the C-terminal domain, far from the mutation site. This observation is consistent with a mutation-induced global effect that influences C-terminal catalytic motifs.

#### CORRELATED motion analysis suggests R48Q experiences global motion reprogramming

To quantify correlated and anticorrelated motions between residue pairs over the course of a trajectory and further illustrate the potential interruption of the motion between the N-terminal and C-terminal regions caused by the R48Q substitution, we performed dynamic cross-correlation matrix (DCCM) analysis (48). By capturing how atomic fluctuations move simultaneously, DCCM provides insight into collective motions, allosteric communication, and dynamic coupling within a protein (49–51). Comparing DCCMs between WT and the R48Q variant enables a direct assessment of how specific mutations alter internal motion patterns, revealing changes in long-range correlations, flexibility, and functional dynamics that may not be apparent from static structures alone. Consistent with our hypothesis, the R48Q variant exhibits a redistribution of motion correlations of regions compared to the WT (**Fig. 3c**). We observed a marked reduction in correlated motion between residues 47-51 in Loop 1 and two segments in the C-terminal domain, including residues 352-360 and 409-422. These three regions are in close contact with one another, and the loss of correlated motion likely reflects disruption of interactions between Loop 1 and its C-terminal contacts. We have also seen a reduction in correlated motion between residues 233-246 and 319-343 at the C-terminal domain. These two segments form helices that lie in close proximity to residues 352-360 and 409-422, which are in contact with Loop 1. The observed changes likely arise from the propagation of reduced motion originating from the disruption at Loop 1. Additionally, the R48Q variant exhibited enhanced coupling across several regions between the N-terminal and C-terminal domains relative to WT, including the pairs 32-38 and 419-424, 134-166 and 320-343, and 91-102 and 373-391. Interestingly, 91-102 includes the key residue Lys94, which is in direct contact with Loop 1 (**Fig. 2c&d**), and 373-391 is the hydrophobic region that accommodates Trp540 (**Fig. 2e, 3a, 3c, & SI Appendix, Fig. S19**). These altered couplings between WT and the R48Q variant suggest the emergence of new dynamic interactions that negatively contribute to activity regulation. These observations highlight that R48Q impacts the internal dynamic network, potentially influencing functional communication residue networks within the protein.

## Discussion

Although bacteria, yeast, archaea, and plants often encode multiple enzymes or isoforms capable of producing L-asparagine, humans rely on a single enzyme, ASNS, to catalyze the ATP-dependent conversion of L-aspartate and L-glutamine into L-asparagine (21, 52–55). Our study provides the first detailed molecular analysis of the ASNSD-associated R48Q variant, linking a single N-terminal substitution to both local dynamic disruption and global interdomain decoupling using a combination of biochemical, structural, and computational approaches.

The reduced catalytic activity of the R48Q variant can be directly attributed to severe dysfunction of Loop 1 at the N-terminal active site. Impairment of glutaminase activity limits ammonia production, thereby compromising the downstream C-terminal synthetase reaction. The striking conformational rearrangement of Loop 1 identified in this work is a functional element of ASNS that has long been overlooked. The altered dynamic ensemble of Loop 1, consistently observed across consensus EM maps and MD simulations, highlights that precise fine-tuning of the active-site configuration is essential for proper N-terminal domain function. Three-dimensional variability analysis further show that this loop reorganization is accompanied by increased conformational heterogeneity of the Cys1 side chain, indicating loss of active-site preorganization essential for efficient glutamine hydrolysis. Importantly, this defect occurs without perturbation of the overall fold or ammonia tunnel architecture, consistent with kinetic data showing preserved substrate binding but a markedly reduced population of catalytically competent enzymes.

In addition to local conformational disruption, the glutamine substitution at residue 48 changes the coordination between the N- and C-terminal domains, as evidenced by reduced synthetase activity with ammonia as the nitrogen source, and results in a disproportionately high production of pyrophosphate relative to L-asparagine. Our cryo-EM structures and MD simulations further reveal long-range perturbations and an altered dynamical network within the synthetase domain, as well as the redistribution of correlated motion across the two domains in R48Q. What stands out in our analysis is the changes at Trp540 and its nearby hydrophobic patch (residues 375-385) in the R48Q variant. Interestingly, reduced fluctuations observed in MD simulations of the 375-385 region in R48Q, which accommodates Trp540, are accompanied by increased correlated motion with the 91-101 region near Loop 1. The essential role of this C-terminal region in regulating ASNS activity is also supported by the identification of a Trp540 mutation in individuals with ASNSD (16). We have validated the loss-of-function of the W540A variant *in vitro* (**SI Appendix, Fig. S21**). The significantly decreased enzyme activity of the W540A variant not only confirms the importance of the regions identified by our cryo-EM data and MD simulations but also implicate a mechanism for the reduced synthetase activity in R48Q, in which the impaired C-terminal activity observed in the R48Q variant is associated with remodeled contact between Loop 1 and Lys94 within the 91-101 region, and that the resulting dynamical correlation alters the dynamics of the 375-385 region, thereby impacting Trp540 and C-terminal tail motion, leading to the decrease in synthetase activity. The decoupling likely underlies the observed disruption in product stoichiometry and highlights the role of coordinated interdomain communication in ASNS catalysis. This model can also explain that the C-terminal tail is indispensable for synthetase activity in the *E. coli* homolog (16, 18, 38). Based on these observations, we propose that the glutamine substitution alters the structural ensemble of Loop 1 to a catalytically incompetent state and further induces global changes in protein motion at the C-terminal domain. Such conformational reprogramming could occur through residue contacts or through changes in the dynamics of the bound water.

Extending our analysis beyond ASNS, we find Arg48 and Loop 1 are conserved within the Class II glutamine amidotransferase (GATase) family, which includes many enzymes in essential biological pathways, such as amidophosphoribosyltransferase, glutamine-fructose-6-phosphate aminotransferase, and L-glutamate synthase (56–59). A sequence alignment of twelve representative GATases revealed that both the flexible Loop 1 and the corresponding arginine residue (analogous to Arg48) are highly conserved across prokaryotic and eukaryotic species (**SI Appendix, Fig. S22**) (32, 60–68). Structural superimposition of apo-GATase structures yielded an average RMSD of 2.05 Å, indicating conserved loop positioning. Remarkably, when comparing the ligand-bound states of the same group of GATases, the RMSD of Loop 1 decreased to 0.54 Å, suggesting a ligand-induced ordering of Loop 1. These data point to a generalizable mechanism in which the conserved arginine stabilizes Loop 1 via hydrogen bonding and supports substrate engagement through direct interactions, a mechanism shared by Class II GATases and critical for catalysis.

Our work also demonstrates the importance of combining biochemical, biophysical, and computational approaches to elucidate the multifaceted impact of disease-linked mutations through the lens of conformational dynamics. Although the chemical mechanism of ASNS has been proposed for other ASNS homologs, understanding the loss-of-function variant underlying ASNSD remains challenging and requires an integrated view of ASNS’s kinetic, structural, and dynamic features. Similarly, pathogenic variant predicting algorithms, such as AlphaMissense (69, 70), flagged R48Q and several key residues identified in this work as potentially pathogenic, experimental approaches remain essential for elucidating the mechanisms underlying the structural and functional disruptions. (**SI Appendix, Fig. S23**).

Importantly, the clinical relevance of these mechanistic insights extends beyond the molecular level. We show that loss of ASNS activity directly reduces L-asparagine availability, which likely contributes to the severe neurological manifestations seen in ASNSD patients.(7, 15, 71) L-asparagine cannot be simply supplemented to compensate for ASNS deficiency in some cases; this might be due to the limited permeability of the blood-brain barrier and the bidirectional nature of amino acid transporters (72, 73). Additionally, sodium-dependent efflux transporters further restrict brain L-asparagine accumulation (73). Moreover, perturbations in ASNS may disrupt the broader amino acid pool, leading to downstream metabolic imbalances that potentially exacerbate disease pathology (74, 75). Therefore, a systematic application of our integrated approach to other ASNSD-linked variants could reveal whether similar patterns of local and global dysfunction exist across the enzyme. Since multiple reported mutations cluster in the N- and C-terminal domains within the dynamic areas identified in this work, this raises intriguing questions about whether they might reciprocally disrupt N-terminal function through the dynamical network or follow distinct mechanistic pathways. Mapping these relationships will be critical for elucidating the shared and divergent mechanisms underlying ASNSD pathogenesis. By expanding these approaches to other pathogenic variants, we can begin to build a comprehensive picture of ASNS dysfunction, ultimately informing the development of targeted interventions for this devastating disorder.

## Materials and Methods

Recombinant human asparagine synthetase (ASNS) WT and variants were expressed in Sf9 cells using a baculovirus system and purified by Ni-NTA affinity chromatography as described previously and in the SI Appendix (23, 30). Protein purity and concentration were assessed by SDS-PAGE and Bradford assay. Protein secondary structure and thermal stability were evaluated by circular dichroism spectroscopy. Enzymatic activity was measured using continuous UV-visible pyrophosphate assays and HPLC-based end-point assays under defined substrate conditions (23, 24, 30). Kinetic parameters were obtained by nonlinear regression analysis.

Cryo-EM samples were prepared by plunge-freezing purified apo-WT and R48Q ASNS on UltrAuFoil grids. Data were collected on a Titan Krios G4 microscope and processed using cryoSPARC (76), yielding the maps at 2.78 Å resolution for apo-WT and 2.81 Å resolution for the R48Q variant. Atomic models were built and refined using Phenix and Coot (**SI Appendix, Fig. S5-S10 & Table S2**) (77). Conformational heterogeneity was analyzed by 3DVA followed by variability refinement as described previously (30).

Molecular dynamics simulations of WT and R48Q ASNS were performed using explicit solvent models (30) and the OPLS_2005 force field. Correlated motions were analyzed using dynamic cross-correlation matrices.

Full experimental procedures, data processing workflows, and computational methods are provided in the SI Appendix.

## Data availability

Primary data are included in the article and/or Supplementary Information. The cryo-EM-derived enzyme structure model has been deposited in the PDB (10LS & 10LT), and the Electron Microscopy Data Bank (EMD-75273 & EMD-75275).

## Acknowledgments and funding source

We thank Dr. Anthony T. Iavarone (University of California, Berkeley, QB3 Mass Spectrometry Facility) and Dr. Peter Randolph (Florida State University, Institute of Molecular Biophysics Protein Biophysics Facility) for their support with mass spectrometry and circular dichroism measurement, respectively. We also thank Ms. Yvonne Lin for her contributions to the clinical literature search and acknowledge the technical support of Drs. Nebojsa Bogdanovic and Jiawei Li at the Southeastern Center for Microscopy of Macromolecular Machines (NIH-R24GM145964). We further acknowledge the Indiana University School of Medicine Electron Microscopy Facility and the Purdue University Electron Microscopy Facility (RRID:SCR_025545), and thank Drs. Vago and Klose for assistance with data collection. This work was supported by the National Institutes of Health (R35GM160337 to W.Z.; S10 OD028723, R01GM111695 to Y.T.), the American Cancer Society (DBG-23-1038947-01-IBCD to Y.T.), the Indiana University School of Medicine (Y.T.), and the Florida State University Start-up Fund (W.Z.). A.C. acknowledges support from the 2025 Fondazione Umberto Veronesi Postdoctoral Fellowship.

