## SupplementaryInfo for "Conformational Remodeling Underlies Activity Loss in Disease-Linked Asparagine Synthetase Variant"

This PDF file includes:

Materials and Methods

Figures S1 to S23

Tables S1 to S2

SI References

### Materials and Methods

#### Recombinant ASNS protein expression and purification

The C-terminally $His_{10}$-tagged human ASNS WT and variants were expressed and purified according to the previously published protocol. Briefly, site-directed mutations were introduced into the pFL-hASNS-TEV-His10 vector (pYT1215) (1) by GenScript (Piscataway, NJ), which was then used to generate recombinant baculovirus expressing C-terminally His10-tagged ASNS and variants (2). The expression and purification of WT human ASNS, as well as ASNS variants, were carried out as described previously (3). Briefly, a 1 L culture of Sf9 cells (1.5 x $10^6$ cells/mL) was infected with the baculovirus expressing recombinant human ASNS or variants at an eMOI of 4.0. The infected cells were then incubated at 27 °C for 72 h, harvested by centrifugation, and frozen in liquid $N_2$. The resulting pellet was then stored at -80 °C until use. All purification steps were performed at 4 °C. The cells were lysed by sonication for 4 minutes in a buffer containing 100 mM EPPS, 300 mM NaCl, 50 mM imidazole, and 5 mM tris(2-carboxyethyl) phosphine (TCEP), pH 8. After centrifugation at 12,000 rpm for 45 minutes, the cell lysate supernatant was loaded onto a Ni-NTA affinity column pre-equilibrated with the same buffer. The column was washed with the same buffer before the target proteins were eluted using a buffer containing 100 mM EPPS, 300 mM NaCl, 250 mM imidazole, and 5 mM TCEP, pH 8. Fractions containing human ASNS were identified and pooled using SDS-PAGE analysis. Protein samples were buffer-exchanged into 100 mM EPPS, 150 mM NaCl, 5 mM TCEP, and 20% glycerol, pH 8, using a PD-10 size-exclusion column. The purified enzyme was concentrated using an Amicon Ultra 30 kDa spin filter, and its concentration was determined using a Bradford assay.(4) Aliquots of the ASNS and variants were stored at -80 °C.

#### Circular dichroism (CD)

The ASNS WT and the R48Q variant used in CD measurements were first exchanged into a buffer containing 10 mM potassium phosphate, pH 8, and concentrated to a final concentration of 0.2 µg/µL using an Amicon Ultra 0.5 mL 30 kDa spin filter. The solution in the flow-through was collected to use as a blank control. The CD measurement was performed using a Chirascan VX spectropolarimeter equipped with a Peltier temperature controller. 150 µL of the enzyme sample or the blank control was used for melting temperature determination. Experimental data were collected from 200 to 280 nm at 0.25s intervals with the temperature ramping from 20 ˚C to 80 ˚C at a scan rate of 2 ˚C /min.

#### UV-vis-based pyrophosphate assay

The rate of inorganic pyrophosphate (PPi) production catalyzed by the ASNS R48Q variant was measured spectrophotometrically using EnzChek™ Pyrophosphate Assay Kit in a continuous assay format. In this assay, the reaction (400 µL) was initiated by the addition of enzyme (400 nM) to a reaction mixture containing 2-amino-6-mercapto-7-methylpurine ribonucleoside (0.2 mM), purine nucleoside phosphorylase (1 U/mL), inorganic phosphatase (0.09 U/mL), ATP, L-aspartate, and L-glutamine or ammonia in a buffer containing 100 mM EPPS, 2 mM dithiothreitol (DTT), and 10 mM $MgCl_2$, pH 8. The absorption of the product was tracked every 0.02 min at 360 nm over a 4-minute period at 37 °C. When L-aspartate was varied (0-12.5 mM), the concentration of L-glutamine (20 mM) or $NH_4Cl$ (100 mM) and the concentration of ATP (5 mM) were held constant. Similarly, when either L-glutamine (0-20 mM) or $NH_4Cl$ (0-100 mM) was varied, the L-aspartate (10 mM) and ATP (5 mM) concentrations were held constant. When ATP (0-5 mM) was varied, the L-aspartate (10 mM), L-glutamine (20 mM), or $NH_4Cl$ (100 mM) concentrations were held constant. Data were analyzed using GraphPad Prism.

#### HPLC-based activity assay

An end-point assay for quantifying L-glutamate and L-asparagine production rates was adapted using HPLC (5). In this assay, the reaction (300 µL) was initiated by the addition of 400 nM enzyme to a reaction mix set at 37 ˚C containing 100 mM EPPS, 2 mM DTT, 10 mM $MgCl_2$, L-aspartate (0-12.5 mM), ATP (0-5mM), and Ammonium chloride (0-100 mM) or L-glutamine (0-20 mM), pH 8. When L-aspartate was varied (0-12.5 mM), the concentration of L-glutamine (20 mM) or $NH_4Cl$ (100 mM) and the concentration of ATP (5 mM) were held constant. Similarly, when either glutamine (0-20 mM) or $NH_4Cl$ (0-100 mM) was varied, the L-aspartate (10 mM) and ATP (5 mM) concentrations were held constant. When ATP (0-5 mM) was varied, the L-aspartate (10 mM), glutamine (20 mM) or $NH_4Cl$ (100 mM) concentrations were held constant. After incubating at various intervals (0-30 minutes), a 30 µL aliquot of the reaction mixture was quenched by 15 µL of glacial acetic acid (AcOH). After precipitation of the enzyme, 25.5 µL of 10 M NaOH and 105 µL of 1 M $Na_2CO_3$ (pH 9) were added to adjust the pH of the solution. Following this, 5 µL of 1-fluoro-2,4-dinitrobenzene (DNFB) was diluted in 145 µL of DMSO. 45 µL of the saturated DNFB solution was then added to the buffered reaction mix. The resulting solution was then incubated at 50 ˚C for 45 minutes. The DNFB reaction was then quenched by adding 30 µL AcOH. Samples were filtered prior to HPLC analysis. Chromatographic separation was carried out on an HPLC system equipped with a Hypersil GOLD™ C18 Selectivity column at a flow rate of 0.25 mL/min using a stepwise gradient of mobile phase B (acetonitrile) in mobile phase A (86% formic acid, pH 3.6). The gradient was as follows: 14% B (0–2 min), 17% B (2–12 min), 20% B (12–17 min), 100% B (17–22 min), followed by re-equilibration at 14% B (22–42 min), achieving baseline separation of L-asparagine, L-glutamate, L-aspartate, and L-glutamine. DNFB-derivatized amino acids were detected at 365 nm.The derivatized amino acid was detected at 365 nm. Standard curves were established using the same experiment procedure, except the reaction mixture was substituted with the known concentration of L-glutamine, L-aspartate, L-glutamate, and L-asparagine. Peak areas were converted into concentration using the standard curves corresponding to each amino acid. The rate of L-asparagine production was analyzed using GraphPad Prism.

**Cryo-EM sample preparation and data Collection**

For cryo-EM analysis, the purified enzyme samples, apo-WT and R48Q, were dialyzed against 50 mM Tris-HCl, pH 8.0, containing 200 mM NaCl and 5 mM beta-mercaptoethanol and concentrated to 3.2 mg/mL using a spin column (MilliporeSigma, St. Louis, MO). 300 mesh UltrAuFoil R1.2/1.3 grids were glow-discharged for 1 minute with a current of 15mA in a PELCO easiGlow system before being mounted onto a Mark IV Vitrobot (FEI/Thermo Fisher Scientific). The sample chamber on the Vitrobot was kept at 4 °C with a relative humidity of 100%. 3.0 µl of each recombinant ASNS sample at a concentration of 3.2 mg/ml was applied to the grid, which was then blotted from both sides for 4 seconds with blot force set at 0. After blotting, the grid was rapidly plunge-frozen into a liquid ethane bath cooled by liquid nitrogen.

Cryo-EM grids were initially screened using a 200 kV Glacios transmission electron microscope (FEI/Thermo Fisher Scientific) equipped with a Falcon 4 direct electron detector. Grids exhibiting optimal ice thickness and particle distribution were selected for high-resolution data collection on a 300 kV Titan Krios G4 transmission electron microscope (FEI/Thermo Fisher Scientific) equipped with a K3 direct electron detector and a BioQuantum energy filter (Gatan). Movies were acquired in super-resolution counting mode using EPU at a nominal magnification of 105,000×, corresponding to a physical pixel size of 0.411 Å. Data were collected over a defocus range of −0.8 to −1.8 µm with a total accumulated electron dose of 60 $e^-/Å^2$. During motion correction, movies were binned two-fold, yielding dose-weighted micrographs with a final pixel size of 0.822 Å. Following motion correction and CTF estimation, micrographs were manually inspected and curated based on relative ice thickness and CTF fit quality. As a result, 6,585 micrographs out of 7,886 for the WT dataset and 6,154 micrographs out of 7,942 for the R48Q variant were retained for subsequent image processing.

**Structure determination by CryoSPARC v4.7.0**

The data processing was carried out by CryoSPARC v4.7.0 (6). As indicated in full workflows in **SI Appendix, Fig. S5 & S8**, structure determination for ASNS WT and the R48Q variant followed the same image-processing workflow unless otherwise noted. Template-based particle picking was performed using a reference generated from the apo-WT structure (PDB: 8SUE). As shown in **SI Appendix, Fig. S5 & S8**, the initially selected particles were subjected to 4 rounds of 2D classification to remove poor particles. These particles were then processed through two iterative rounds of *Ab initio* reconstruction followed by heterogeneous refinement, yielding 3 initial 3D maps for WT and 3 maps for the R48Q variant. Particles contributing to the highest-quality map of WT or R48Q variant were selected and subjected to an additional cycle of *Ab initio* reconstruction and heterogeneous refinement, followed by a second iteration of the same procedure, generating three further 3D reconstructions. Particles corresponding to the best map from this stage were refined using non-uniform refinement with C1 symmetry, including CTF and defocus refinement, resulting in maps at 3.0 Å resolution for WT and 3.2 Å resolution for the R48Q variant. Subsequent non-uniform refinement imposing C2 symmetry improved the resolutions to 2.98 Å (WT) and 3.0 Å (R48Q). Particles contributing to these maps were then re-extracted from the micrographs using a box size of 320 pixels. The re-extracted particles underwent an additional round of non-uniform refinement with C1 symmetry, followed by a final round of non-uniform refinement incorporating global CTF, defocus, and beam-tilt corrections with C2 symmetry imposed. This final refinement yielded reconstructions at overall resolutions of 2.78 Å for ASNS WT and 2.81 Å for the R48Q variant, as estimated by Fourier shell correlation (FSC) using the gold-standard 0.143 cut-off criterion (**SI Appendix, Fig. S6, S7, S9, & S10**) (7–9). The final density for WT map was sharpened using DeepEMhancer (10) via the COSMIC2 platform (11). All cryo-EM maps were visualized using UCSF Chimera (12).

**Model building and refinement**

Each EM map was sharpened using the DeepEMhancer tool (10) available through the COSMIC$^2$ science gateway (13), and the resulting map was used for model building. Initial fitting was performed by placing the apo-ASNS EM structure (PDB: 8SUE) into the sharpened EM map using rigid-body refinement in REFMAC . This initial model was further refined through iterative rounds of real-space refinement in Phenix (11), followed by manual inspection and adjustment in Coot. Final model refinement statistics are summarized in **SI Appendix, Table S2**.

**3D variability analysis (3DVA) and refinement of ASNS WT and the R48Q variant**

EM map of WT or R48Q variant was subjected to 3D variability analysis (3DVA) in cryoSPARC v4.7.0 with 5 principle components (14). A total of 20 frames generated by 3DVA described above will be subjected to variability refinement using *Phenix.varref* program as described (15), yielding a total of 20 PDB files corresponding to each frame for WT as well as the R48Q variant.

**Molecular dynamics simulation**

The full-length WT ASNS model was obtained as described in our previous work (3). The R48Q variant was generated by mutating residue Arg48 to glutamine in the WT model. Protonation states of all ionizable residues at pH 7.4 were assigned using the PROPKA algorithm, as implemented in Maestro (Schrödinger Suite 2024-03). Each model was energy-minimized and then solvated in a TIP3P water box with a minimum of 10 Å between any protein atom and the box boundary. Sodium ions were added to neutralize the total system charge. Following solvation, each system was heated to 300 K and equilibrated in the NPT ensemble (P = 101,325 Pa). For both WT and R48Q, four independent 200-ns NPT MD simulations were performed using the OPLS_2005 force field. All simulations employed periodic boundary conditions (cubic box geometry), with long-range electrostatics treated using particle mesh Ewald and short-range nonbonded interactions truncated at 9 Å.

**Dynamic cross-correlation matrix (DCCM)**

Dynamic cross-correlation matrices (DCCM) were calculated from the atomic fluctuations of the $C_\alpha$ atoms over the course of the molecular dynamics simulations. The correlation coefficient between the motions of atoms *i* and *j* is defined as:

$$C_{ij} = \frac{\langle \Delta \mathbf{r}_i \cdot \Delta \mathbf{r}_j \rangle}{\sqrt{\langle \Delta \mathbf{r}_i \hat{\ } 2 \rangle \langle \Delta \mathbf{r}_j^2 \rangle}}$$

where $\Delta r_i$ is the displacement from the mean position of the *i*th atom. The resulting $C_{ij}$ values range from -1 to 1, indicating fully anti-correlated to fully correlated motions, respectively. To capture differences in correlated motions between systems, the delta DCCM ($\Delta C_{ij}$) was calculated as the element-wise difference between the average DCCMs of the systems under comparison. In this study, the DCCMs were first calculated for each replica and subsequently averaged to obtain a representative DCCM per system. The delta DCCM was then computed from these averaged matrices, providing a robust measure of differential correlated motions induced by the experimental condition. DCCM analysis and visualization were performed using the MDAnalysis package and custom in-house Python scripts.

### Figures

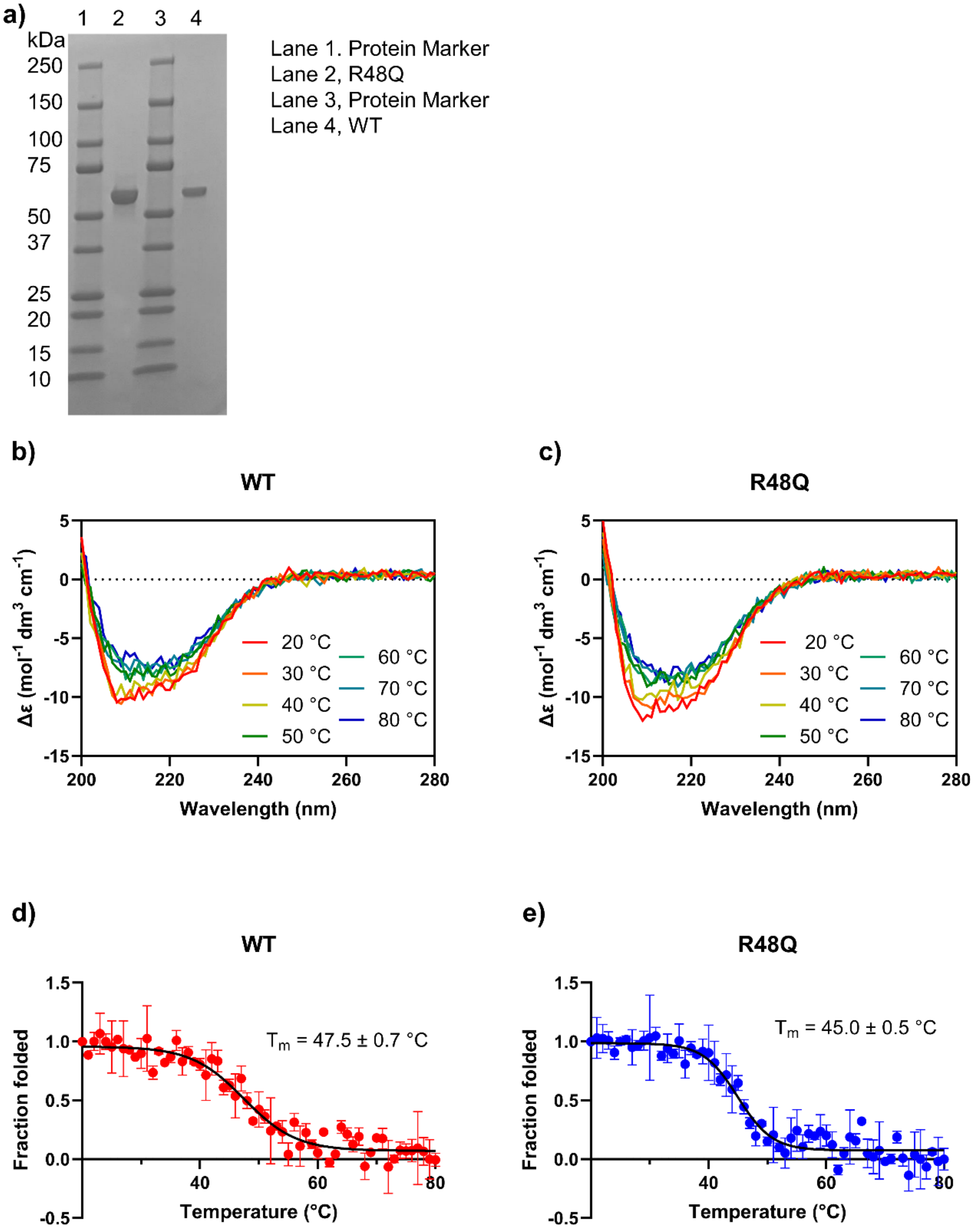


**Fig. S1. Thermal stability of WT and R48Q variants.** a) ASNS WT and R48Q were expressed, purified, and used in this work. b) Far-UV CD spectra recorded from 20 °C to 80 °C reveal temperature-dependent changes in secondary structure for WT. c) Far-UV CD spectra recorded from 20 °C to 80 °C reveal temperature-dependent changes in secondary structure for R48Q. d) Thermal denaturation curves monitored by circular dichroism (CD) at 222 nm of WT with calculated melting temperature. e) Thermal denaturation curves monitored by CD at 222 nm of R48Q with calculated melting temperature.

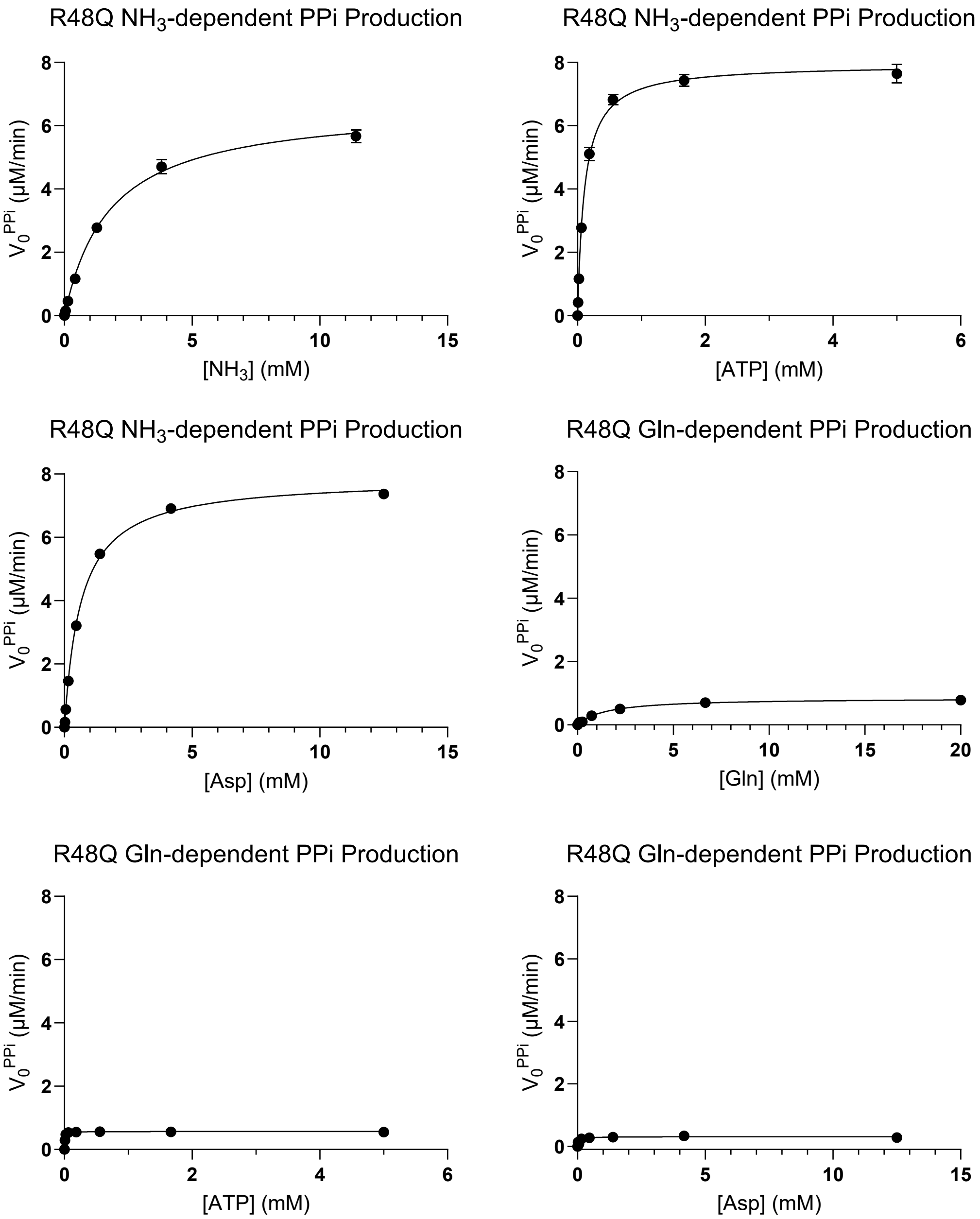


**Fig. S2. Steady-state kinetics of pyrophosphate production in the R48Q-catalyzed reaction.** Michaelis-Menten plots display the initial velocity of pyrophosphate production relative to substrate concentration. Assays were conducted with either glutamine (Gln) or ammonia ($NH_3$) as a nitrogen source. When varying one substrate, the concentrations of the other two remained constant. The R48Q variant shows strong activity with exogenous ammonia, while glutamine-dependent activity is significantly impaired under all conditions. All reactions were performed in duplicate.

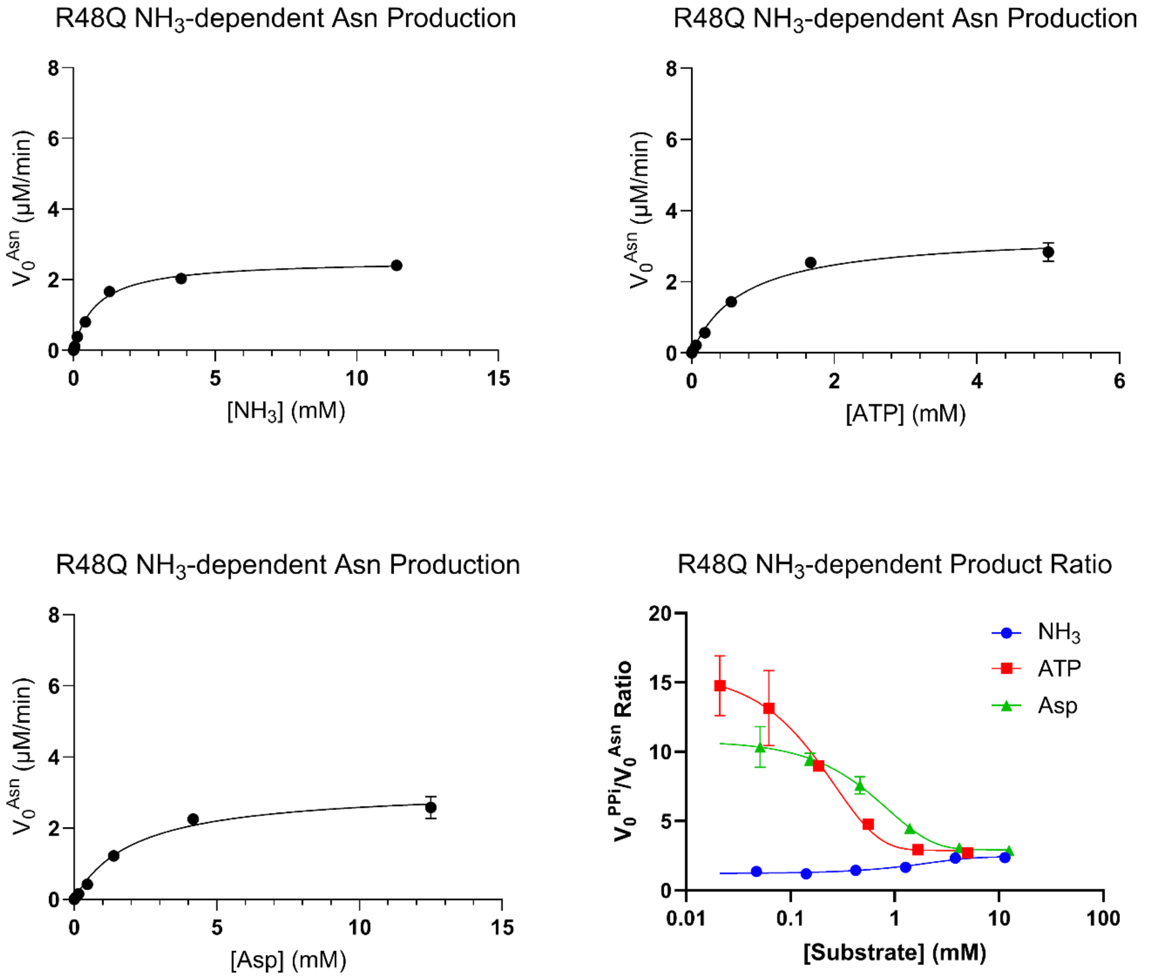


**Fig. S3. Steady-state kinetics of asparagine production in the R48Q-catalyzed reaction.** Michaelis-Menten plots show the initial velocity of asparagine (PPi) production as a function of substrate concentration. Assays were performed in the presence of ammonia ($NH_3$) as a nitrogen source. Under the same concentration of ATP and aspartate, negligible activity was detected when glutamine was used as a substrate. When titrating one substrate, the concentration of the other two substrates remains saturated. All reaction was performed in duplicates. Bottom right: Substrate-dependent modulation of product ratio in R48Q-catalyzed ammonia-dependent reactions. The ratio of pyrophosphate production to asparagine production was determined across varying concentrations of individual substrates (ammonia, blue circles; ATP, red squares; aspartate, green triangles), with the other two substrates held at saturating levels. All reactions were performed in duplicate.

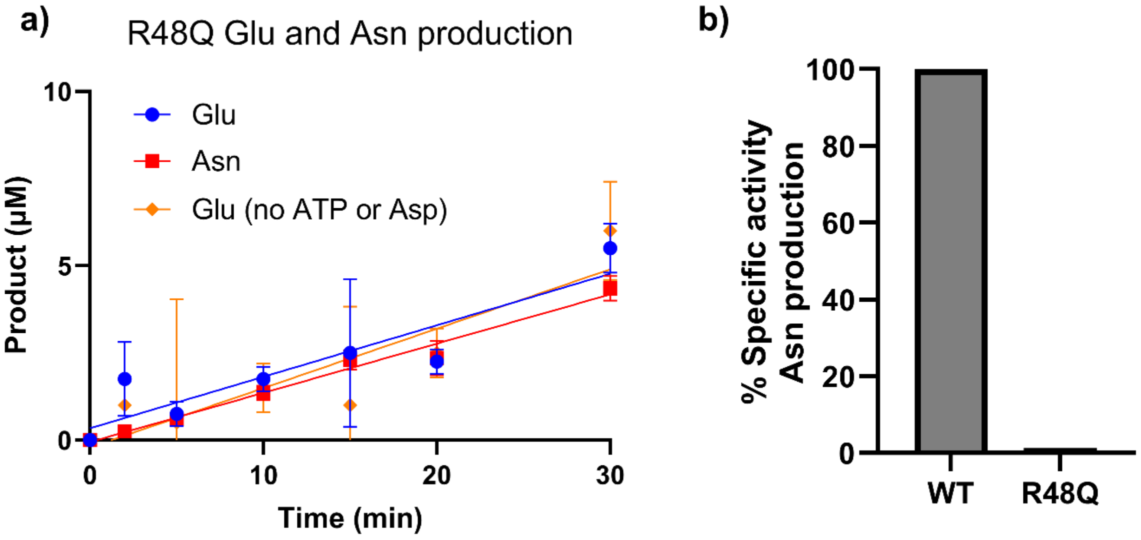


**Fig. S4. Glutamate production of the R48Q variant.** a) Time-course analysis of product formation by the R48Q variant over 30 minutes. Glutamate (Glu, blue circles) and asparagine (Asn, red squares) production were monitored in the presence of all substrates. Glu production was also measured in the absence of ATP and aspartate (Asp, orange diamonds) to assess basal glutaminase activity independent of C-terminal active site engagement. b) Comparison of the specific glutaminase activity between WT and R48Q in the presence of C-terminal substrates. The R48Q variant shows elevated uncoupled glutaminase activity, indicating impaired interdomain communication and loss of catalytic coordination. All reactions were performed in duplicate.

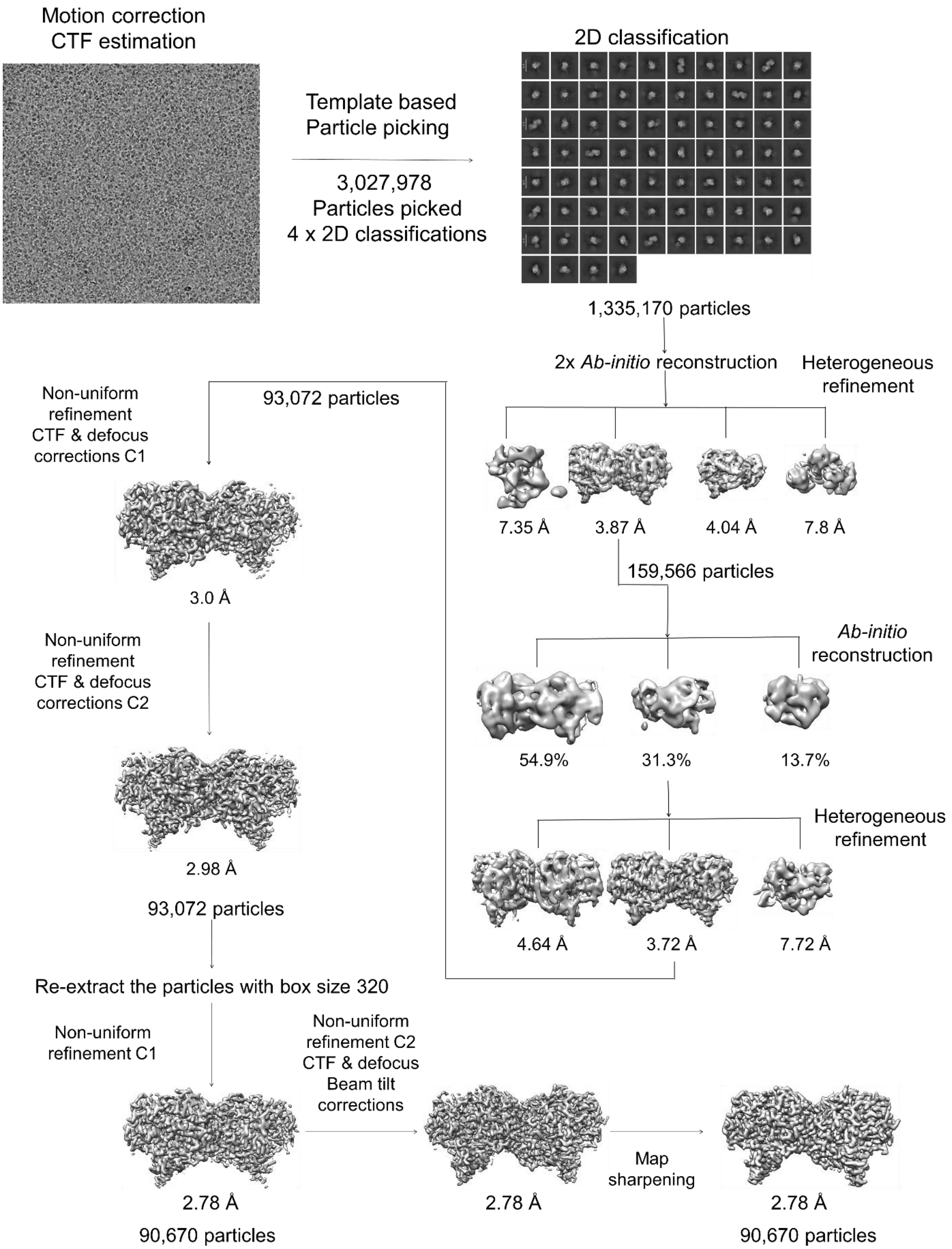


**Fig. S5. WT cryo-EM data analysis workflow.** Summary of the cryo-EM data processing workflow of WT ASNS using cryoSPARC v4.7.0.

### a) Gold standard FSC curve for ASNS (WT)

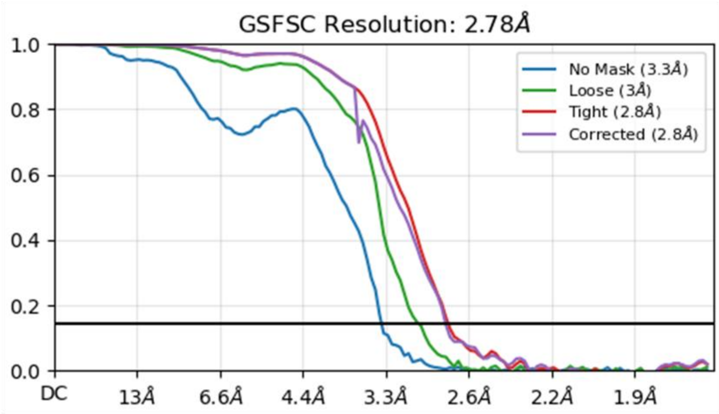


### b) Viewing distribution plots

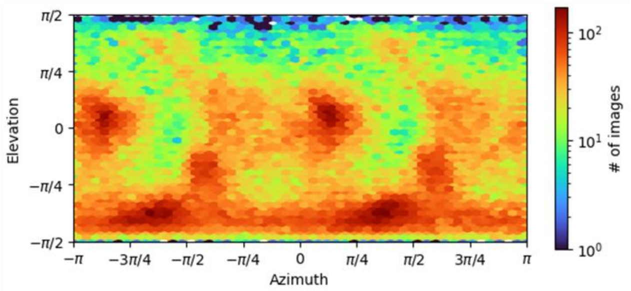


### c) cFSC and cFAR

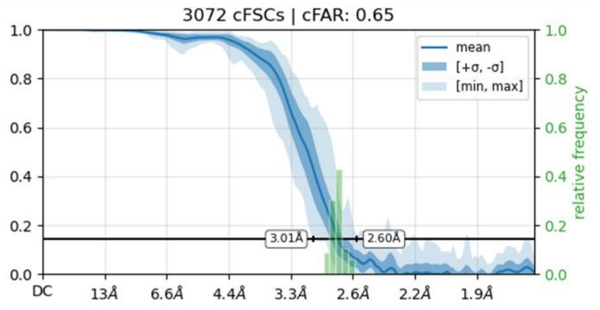


### d) Local resolution

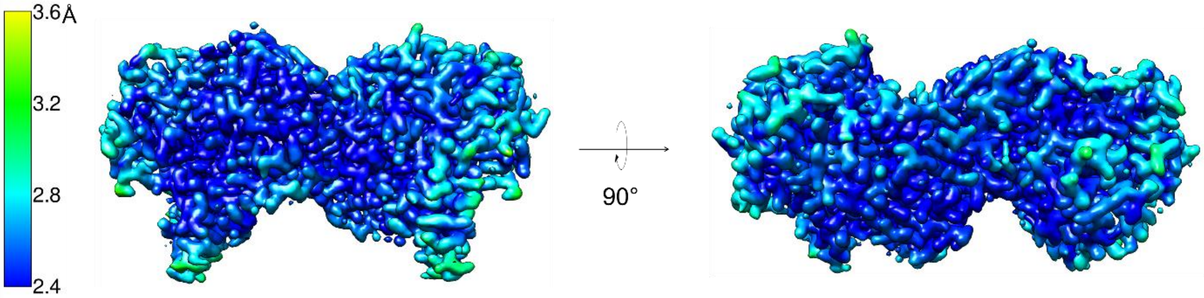


**Fig. S6. Assessment of the quality of the cryo-EM map of WT ASNS.** a) Gold standard FSC curve for the map of ASNS (WT) complex generated in cryoSPARC v4.7.0. b) Viewing distribution plots for the map generated in cryoSPARC v4.7.0. c) Measure of anisotropy is indicated by cFSC and Cfar. d) Local resolution of the map (color scale shown on the left).

### a) Representative densities and corresponding models for ASNA (WT)

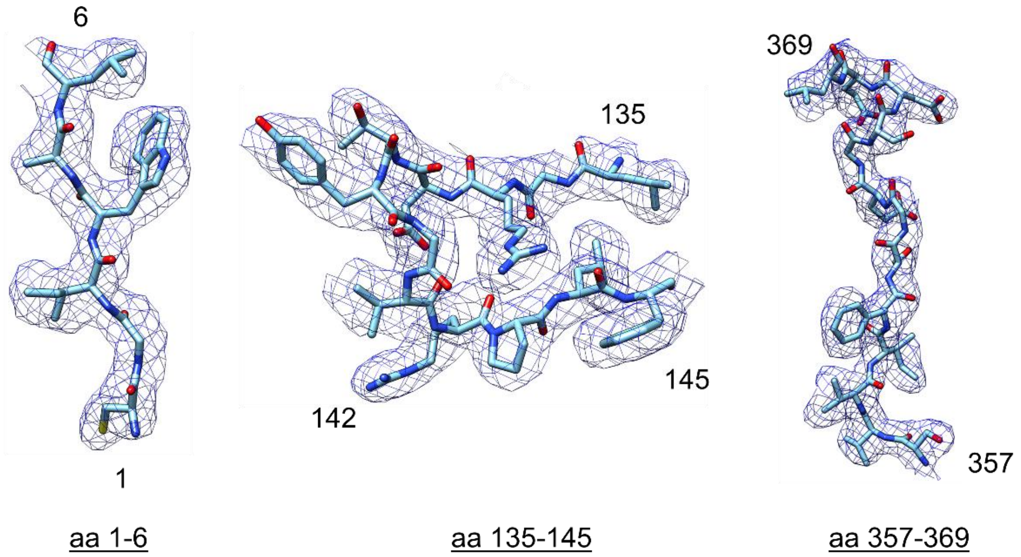


### b) Map-model FSC curve

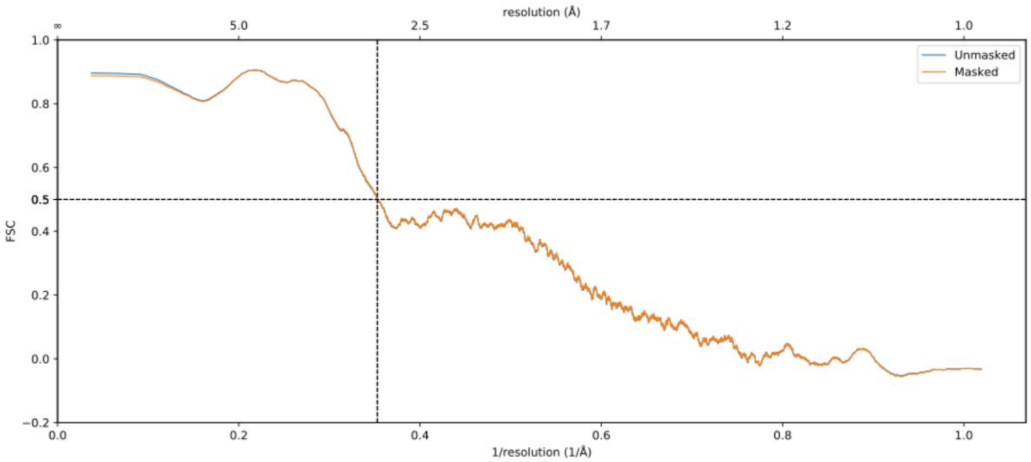


**Fig. S7. Validating the cryo-EM structure: map and WT ASNS model.** a) Representative density of the map for residues in the N-terminal catalytic region (aa1-6), the ammonia tunnel region (aa 135-145), and the C-terminal active site (aa 357-369). b) Model-to-map FSC curves generated in Phenix.

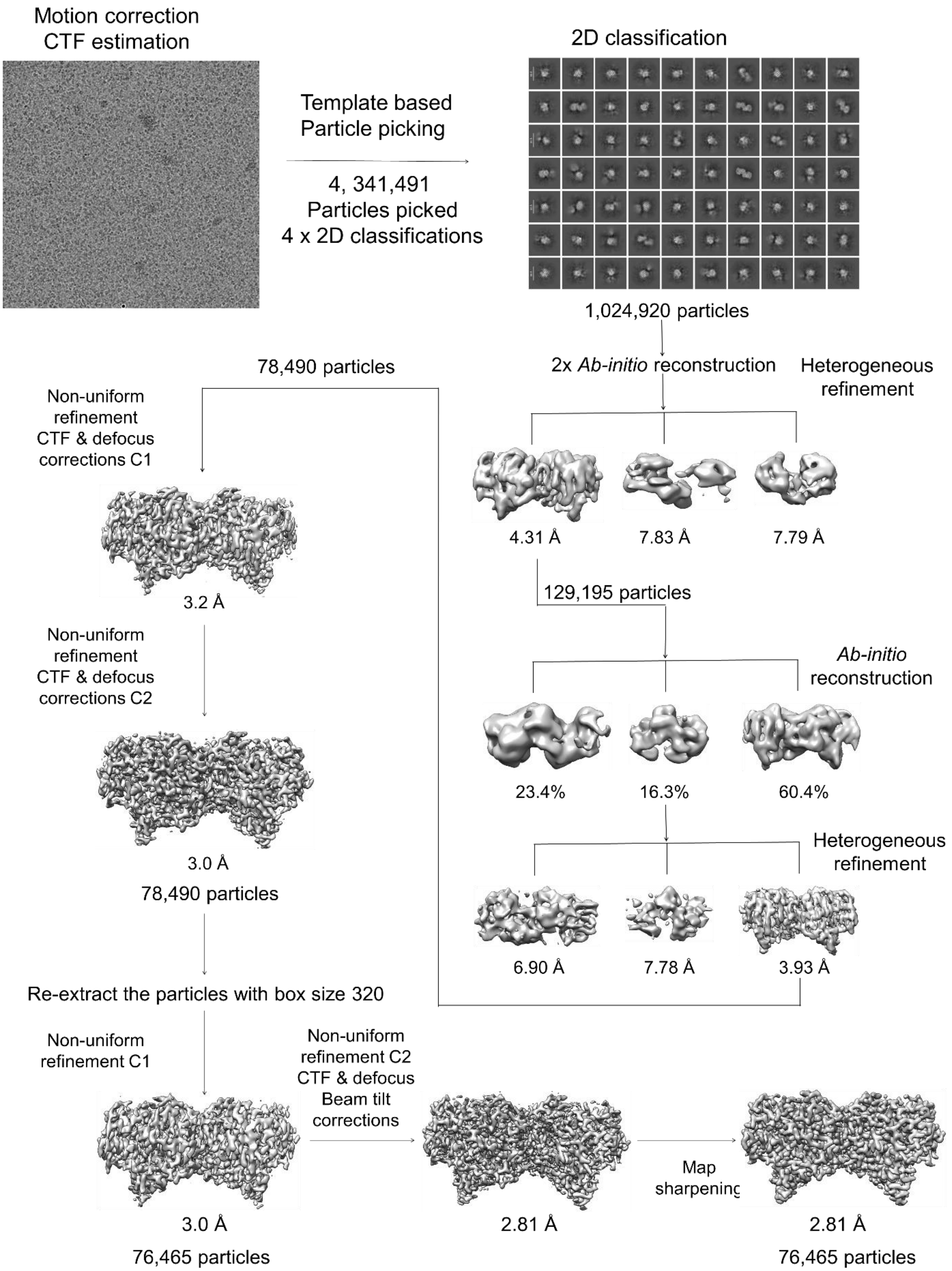


**Fig. S8. Summary of the cryo-EM data processing workflow of the R48Q variant cryo-EM dataset using cryoSPARC v4.7.0.**

### a) Gold standard FSC curve for ASNS (R48Q)

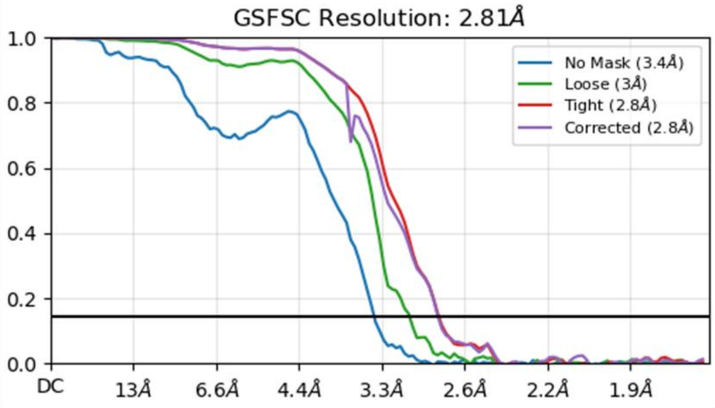


### b) Viewing distribution plots

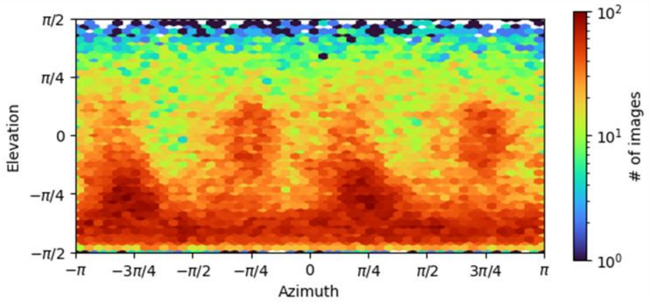


### c) cFSC and cFAR

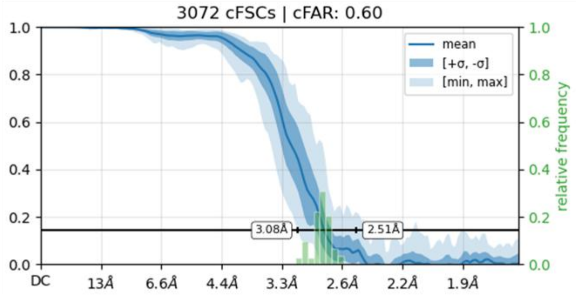


### d) Local resolution

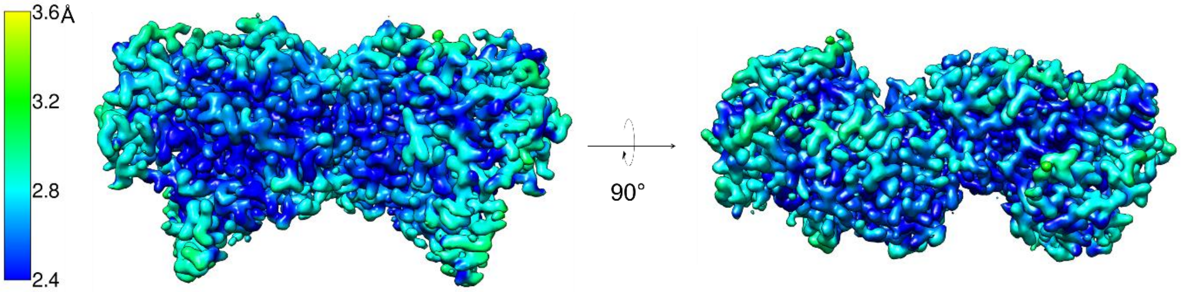


**Fig. S9. Assessment of the quality of cryo-EM map of the R48Q variant.** a) Gold standard FSC curve for the map of ASNS (WT) complex generated in cryoSPARC v4.7.0. b) Viewing distribution plots for the map generated in cryoSPARC v4.7.0. c) Measure of anisotropy is indicated by cFSC and cFAR. d) Local resolution of the map (color scale shown on the left).

### a) Representative densities and corresponding models for ASNA (R48Q)

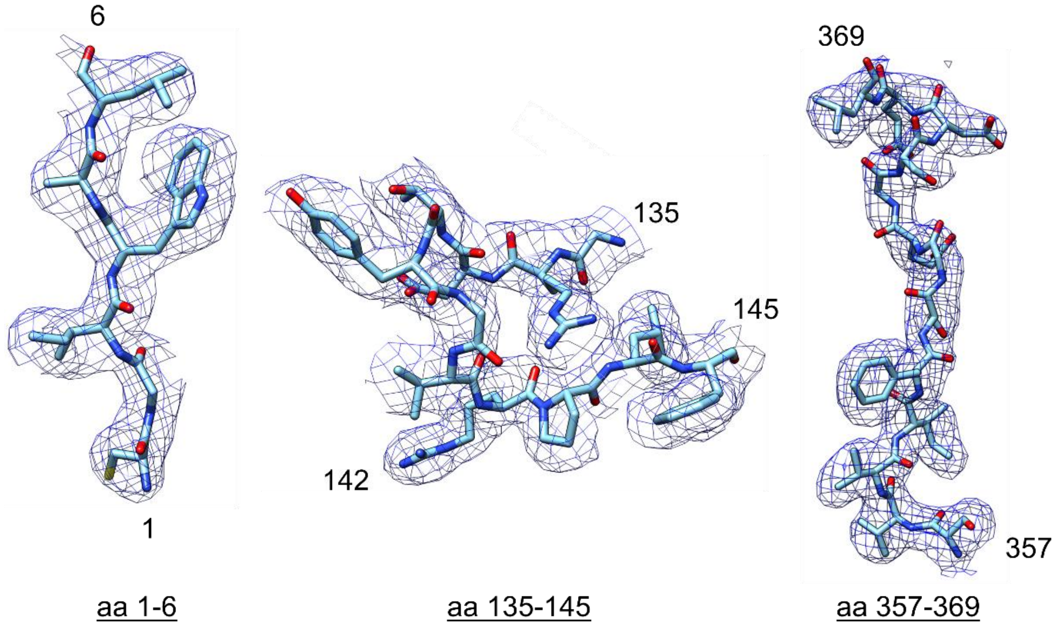


### b) Map-model FSC curve

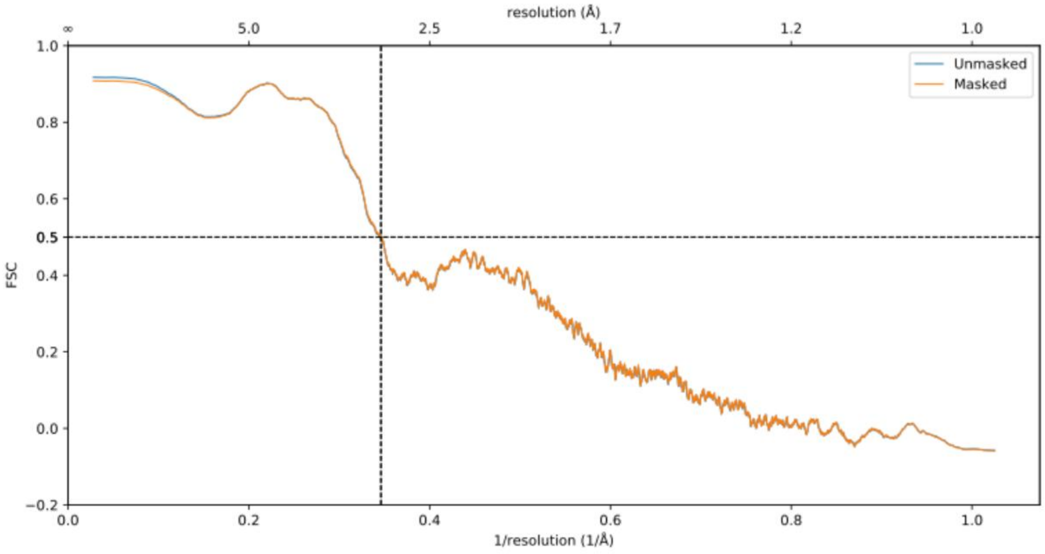


**Fig. S10. Validating the cryo-EM structure: map and its model of the R48Q variant.** a) Representative density of the map for residues in the N-terminal catalytic region (aa1-6), the ammonia tunnel region (aa 135-145), and the C-terminal active site (aa 357-369). b) Model-to-map FSC curves generated in Phenix.

**a)**

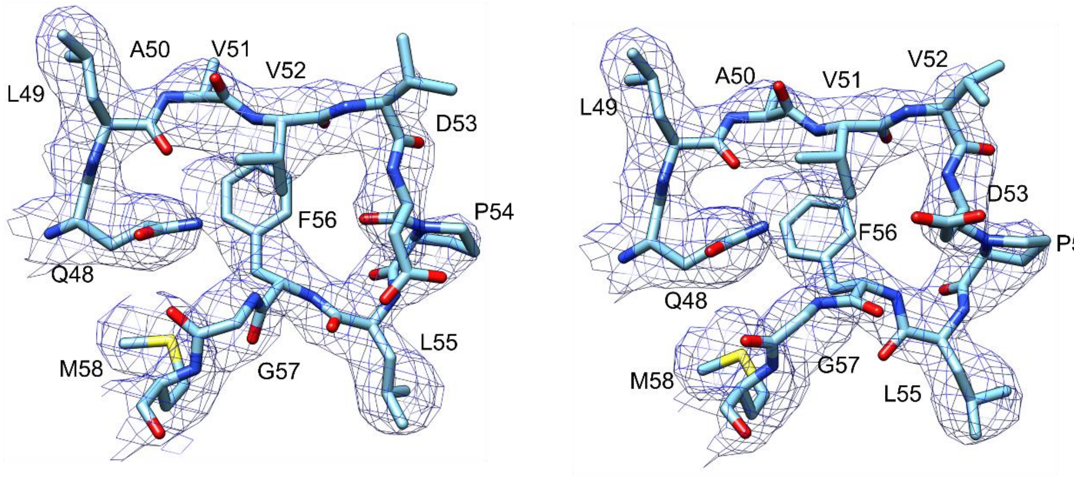


Apo-R48Q Loop 1 chain A    Apo-R48Q Loop 1 chain B

**b)**

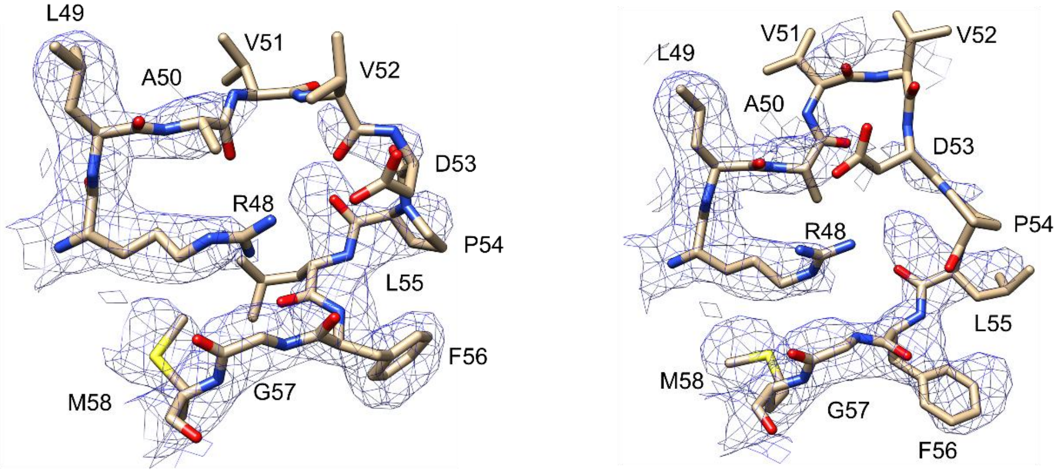


Apo-WT Loop 1 chain A    Apo-WT Loop 1 chain B

**Fig. S11. Cryo-EM maps and corresponding atomic models for Loop 1.** a) Close-up views of Loop 1 in the apo-R48Q variant, comprising residues Q48-M58 (Q48, L49, A50, V51, V52, D53, P54, L55, F56, G57, M58). Loop 1 residues are shown in sky blue, with the corresponding cryo-EM density displayed in blue mesh. The model and map for Chain A are shown on the left, and those for Chain B are shown on the right. b) Close-up views of Loop 1 in the apo-WT ASNS, comprising residues R48-M58 (R48, L49, A50, V51, V52, D53, P54, L55, F56, G57, M58). Loop 1 residues are shown in goldenrod, with the corresponding cryo-EM density displayed in blue mesh. The model and map for Chain A are shown on the left, and those for Chain B are shown on the right.

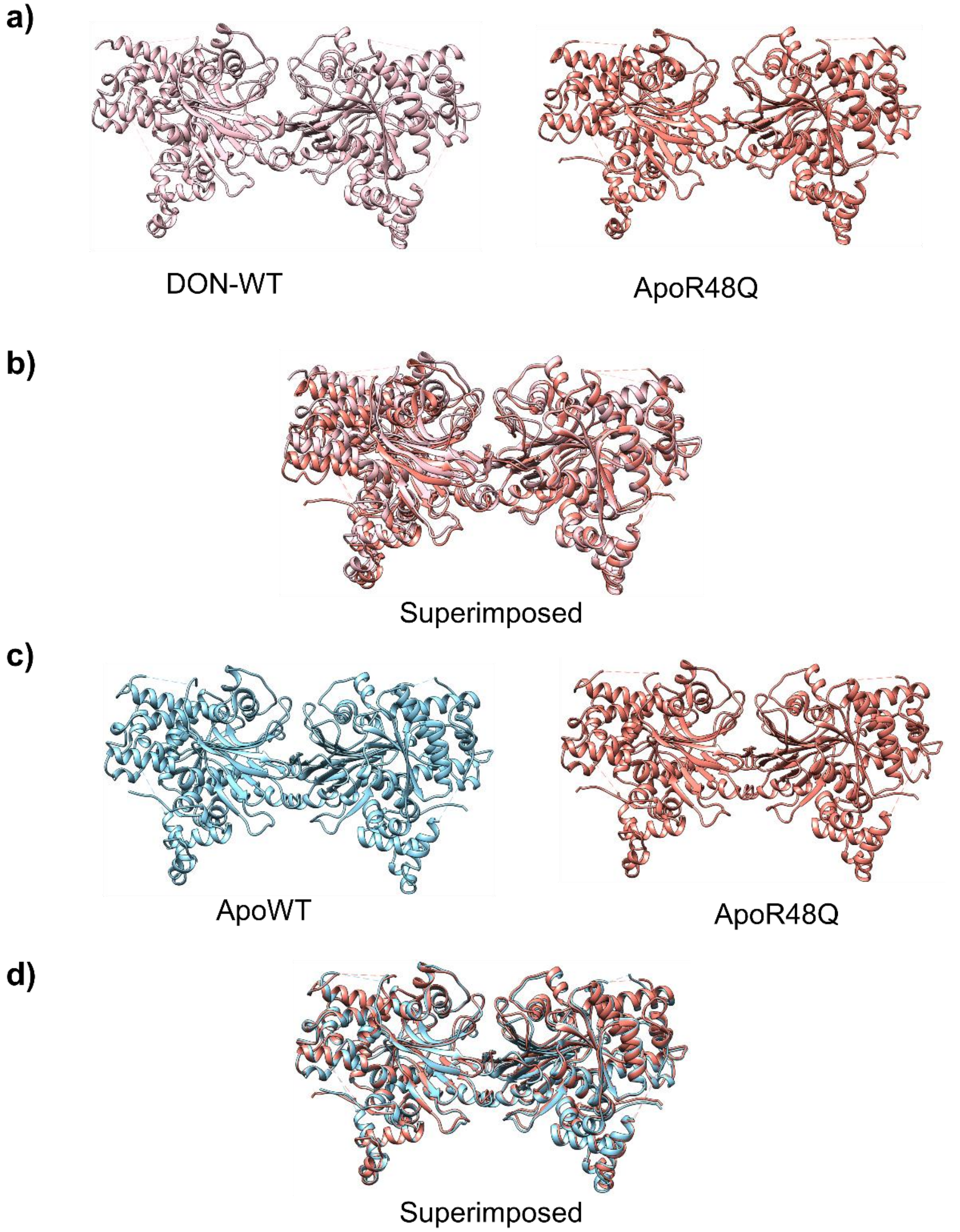


**Fig. S12. Structure of R48Q ASNS and its comparison to DON-WT and apo-WT ASNS.** a) Comparison between DON-ASNS (PDB: 6GQ3 colored in pink) vs. R48Q variant (PDB: 10LT colored in salmon) structures. b) Superimposed models of DON-ASNS and R48Q variant (RMSD: 0.596 Å). c) Comparison between apo-ASNS (PDB: 10LS colored in sky blue) vs. R48Q variant structures. d) Superimposed models of apo-ASNS and the R48Q variant (RMSD: 0.392 Å)

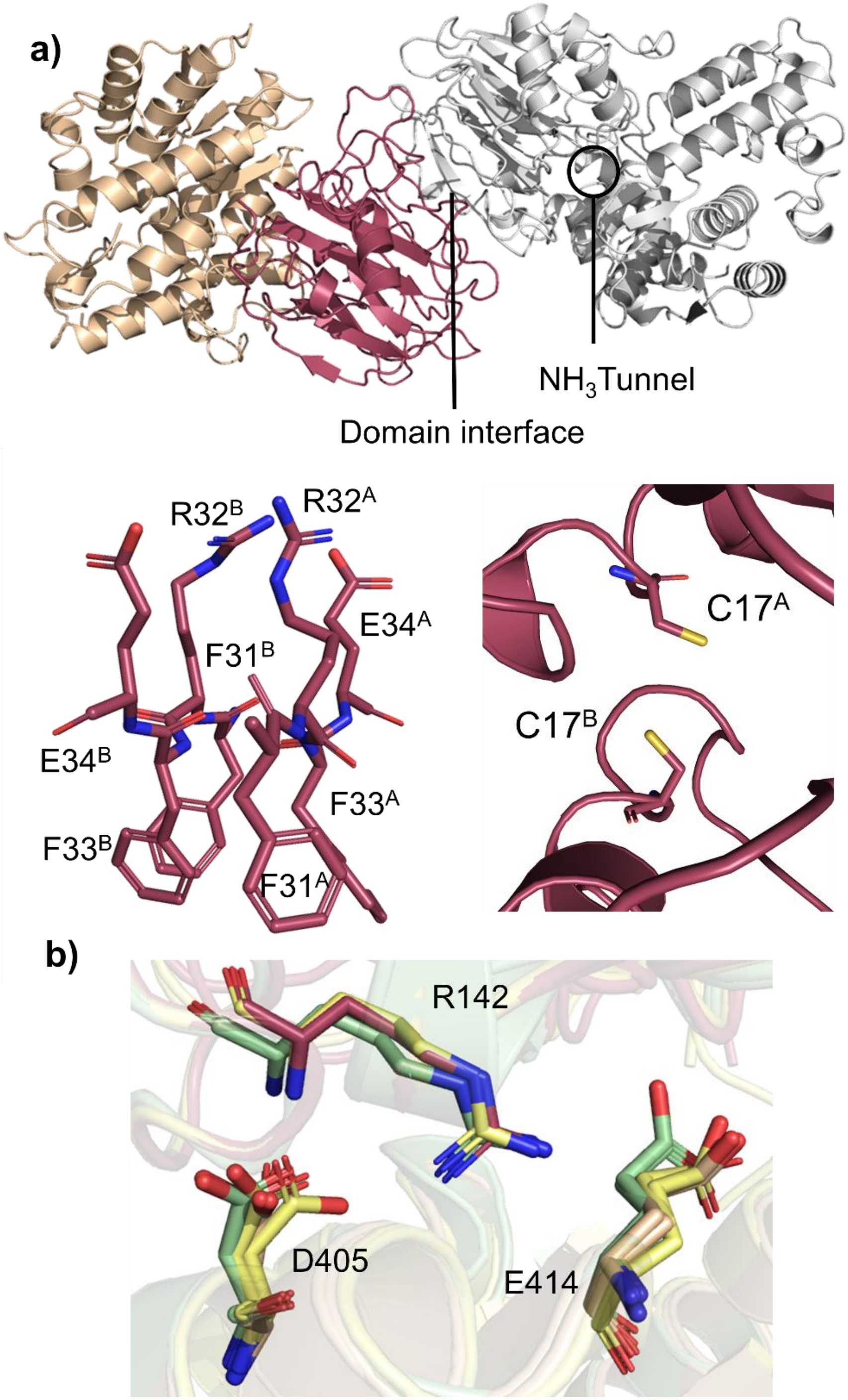


**Fig. S13. R48Q does not alter the overall fold of ASNS.** a) Overall structure of the R48Q ASNS homodimer, with the N-terminal domain in garnet and the C-terminal domain in wheat. A close-up view of the dimer interface highlights stabilizing salt bridges and hydrophobic contacts. The Cys17 residues from chains A and B do not form a disulfide bond in the R48Q structure. b) Structural comparison of interactions near the tunnel-lining residue Arg142 in both monomers across apo R48Q (wheat), apo WT (yellow), and DON-modified WT ASNS (green). Chain A and B are superimposed, and key residues are highlighted in sticks.

a)

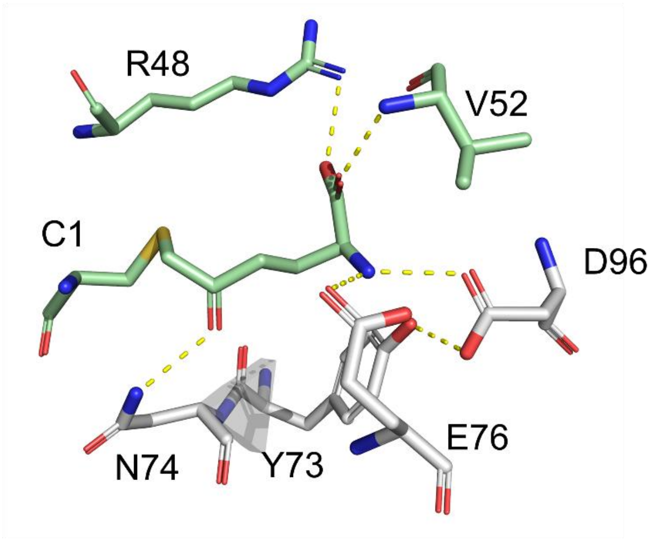


**DON-WT**

b)

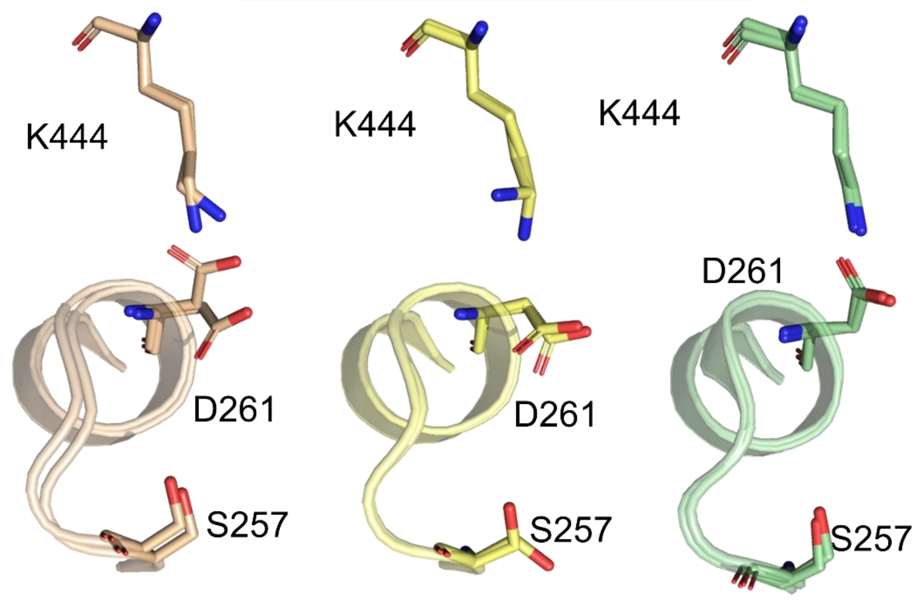


**R48Q** **Apo-WT** **DON-WT**

**Fig. S14. Structural comparison between WT and R48Q.** a) Arg48 interacts with Val52 in Loop 1 and DON in the DON-WT structure (green). Additional interactions between DON and the N-terminal residues in DON-WT are highlighted in gray. g) PP-motif at the C-terminal active site in R48Q (wheat), apo-WT (yellow), and DON-modified WT ASNS (green).

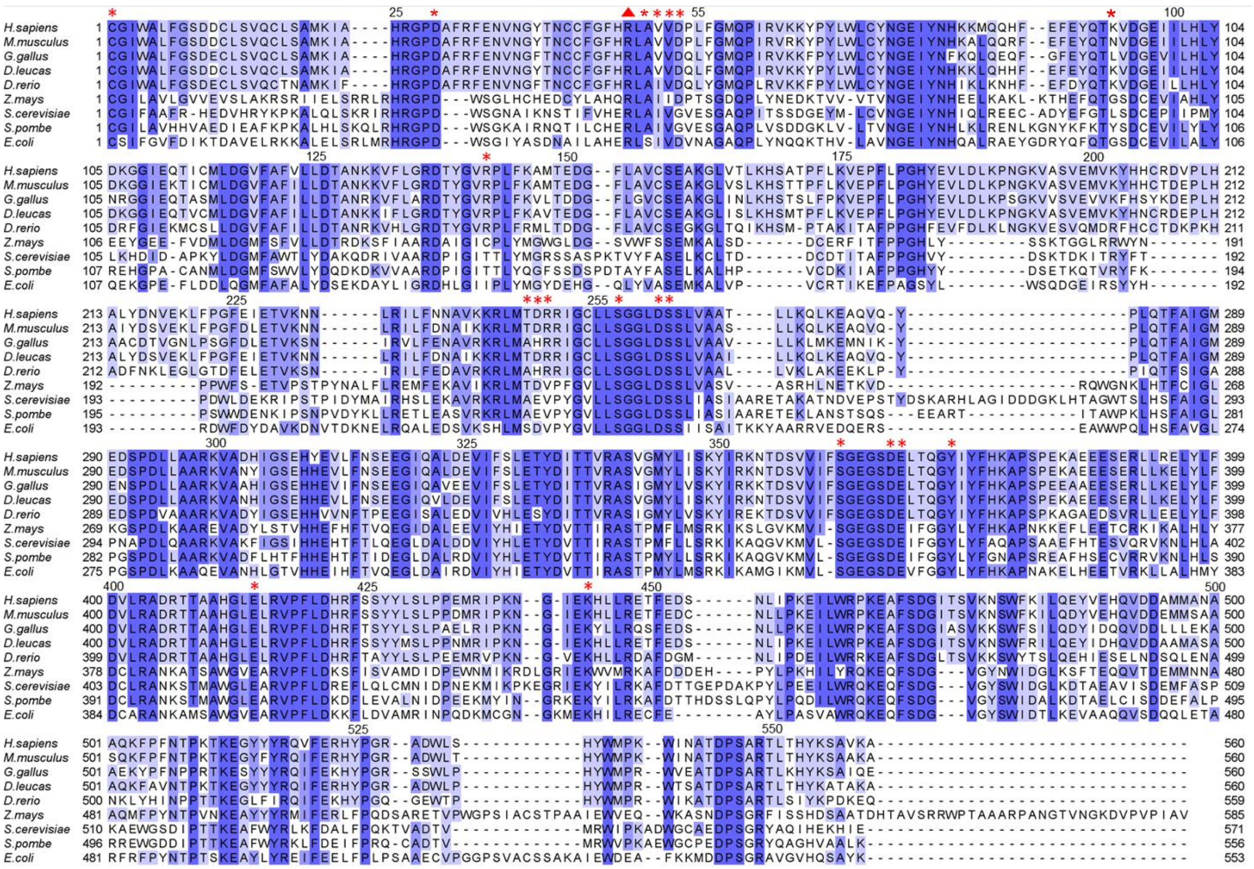

**Fig. S15. Sequence alignment of asparagine synthetases from various organisms.** Conserved residues are highlighted in blue, with color intensity indicating the degree of conservation. The conserved arginine residue in Loop 1 is marked with a red triangle. Red asterisks denote key interacting residues in the apo-WT and DON-WT structures.

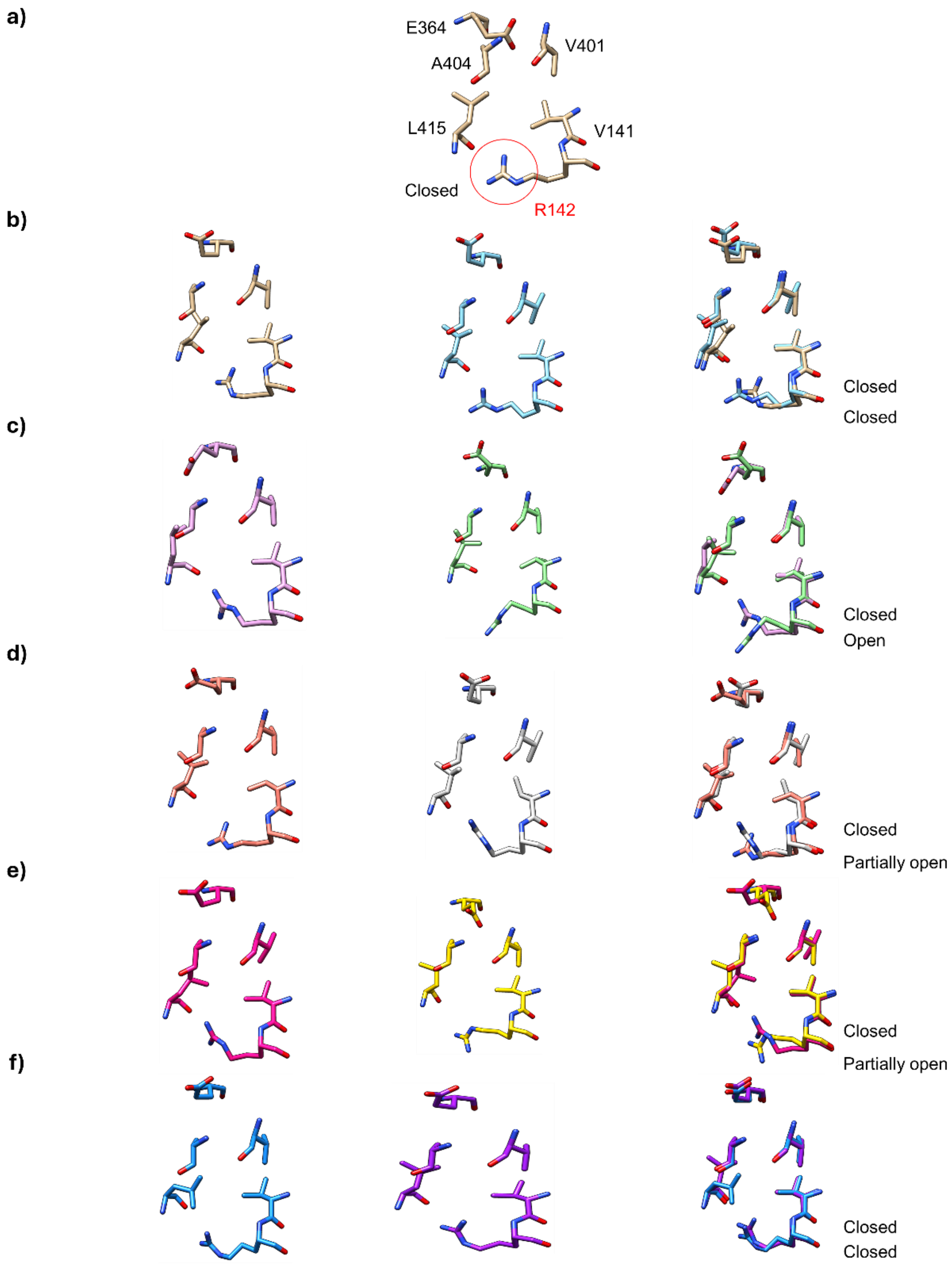


**Fig. S16. PCA-derived conformations of the ammonia tunnel in WT ASNS obtained from 3DVA.** a) Side view of key residues (Val141, Arg142, Glu364, Val401, Ala404, Val414, Leu415) forming the ammonia tunnel in the EM map of apo-ASNS. The side chain of Arg142, which blocks the tunnel, is highlighted (red circle). (b–f) Side views of the same residues in variable EM maps from 3DVA of WT, with corresponding models generated by 3D variability refinement. Residues from the tunnel conformations at frame 0 (one end) and frame 19 (the opposite end) are superimposed for each PCA component: b) component 0, c) component 1, d) component 2, e) component 3, f) component 4.

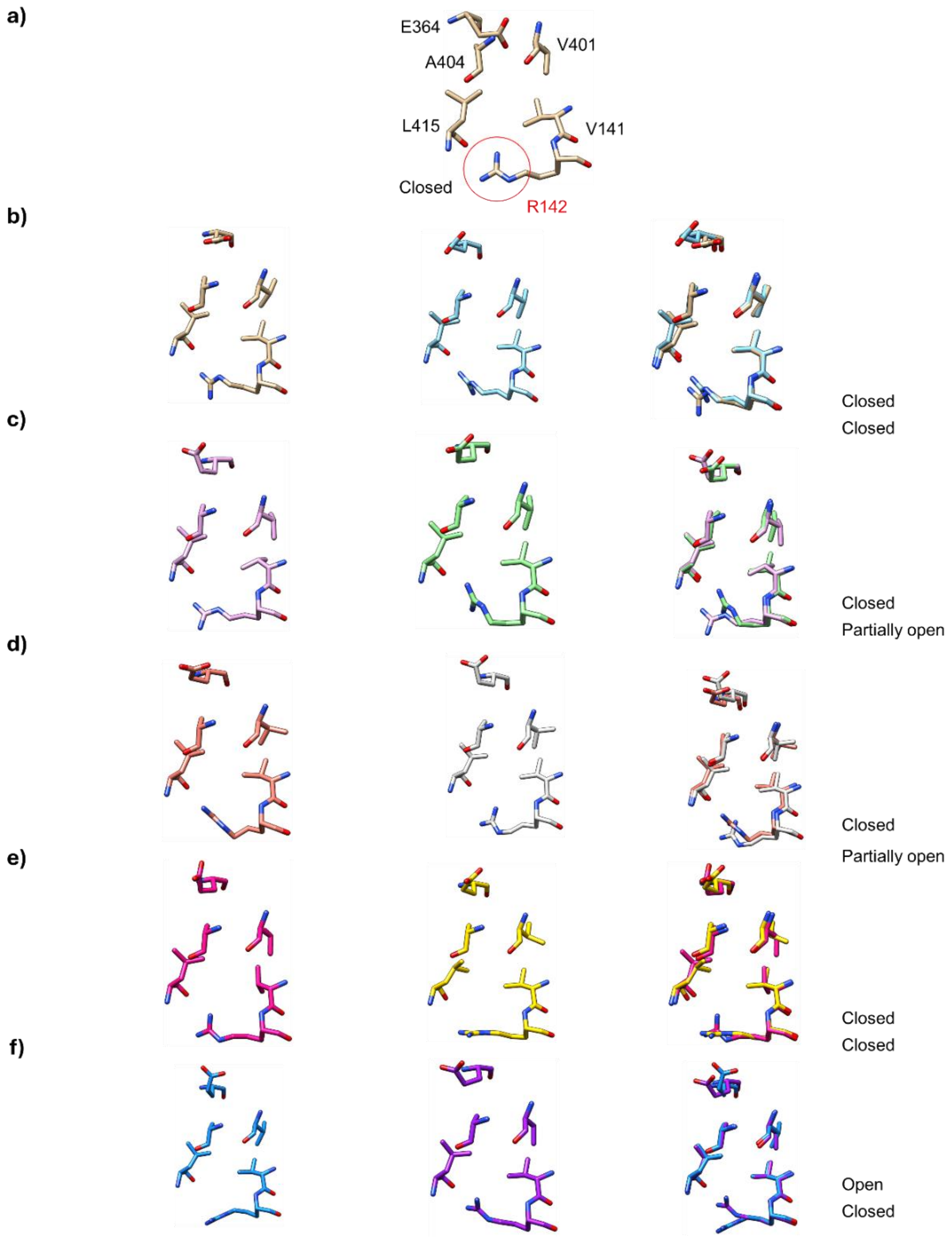


**Fig. S17. PCA-derived conformations of the ammonia tunnel in the R48Q variant obtained from 3DVA.** a) Side view of key residues (Val141, Arg142, Glu364, Val401, Ala404, Val414, Leu415) forming the ammonia tunnel in the EM map of R48Q. The side chain of Arg142, which blocks the tunnel, is highlighted (red circle). (b–f) Side views of the same residues in variable EM maps from 3DVA of the ASNS (R48Q), with corresponding models generated by 3D variability refinement. Residues from the tunnel conformations at frame 0 (one end) and frame 19 (the opposite end) are superimposed for each PCA component: b) component 0, c) component 1, d) component 2, e) component 3, f) component 4.

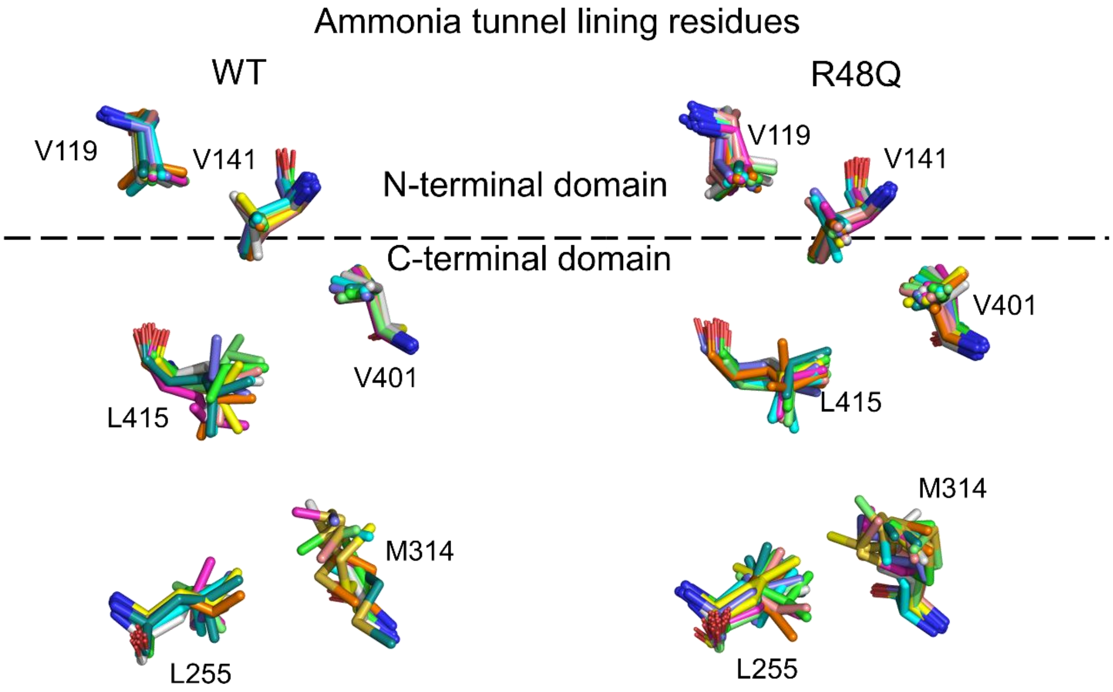


**Fig. S18. 3DVA analysis of the tunnel lining residues in the R48Q variant.** Tunnel lining residues in the representative 3DVA-derived structures, including Val119, Val141, Leu255, Met344, Val401, and Leu415.

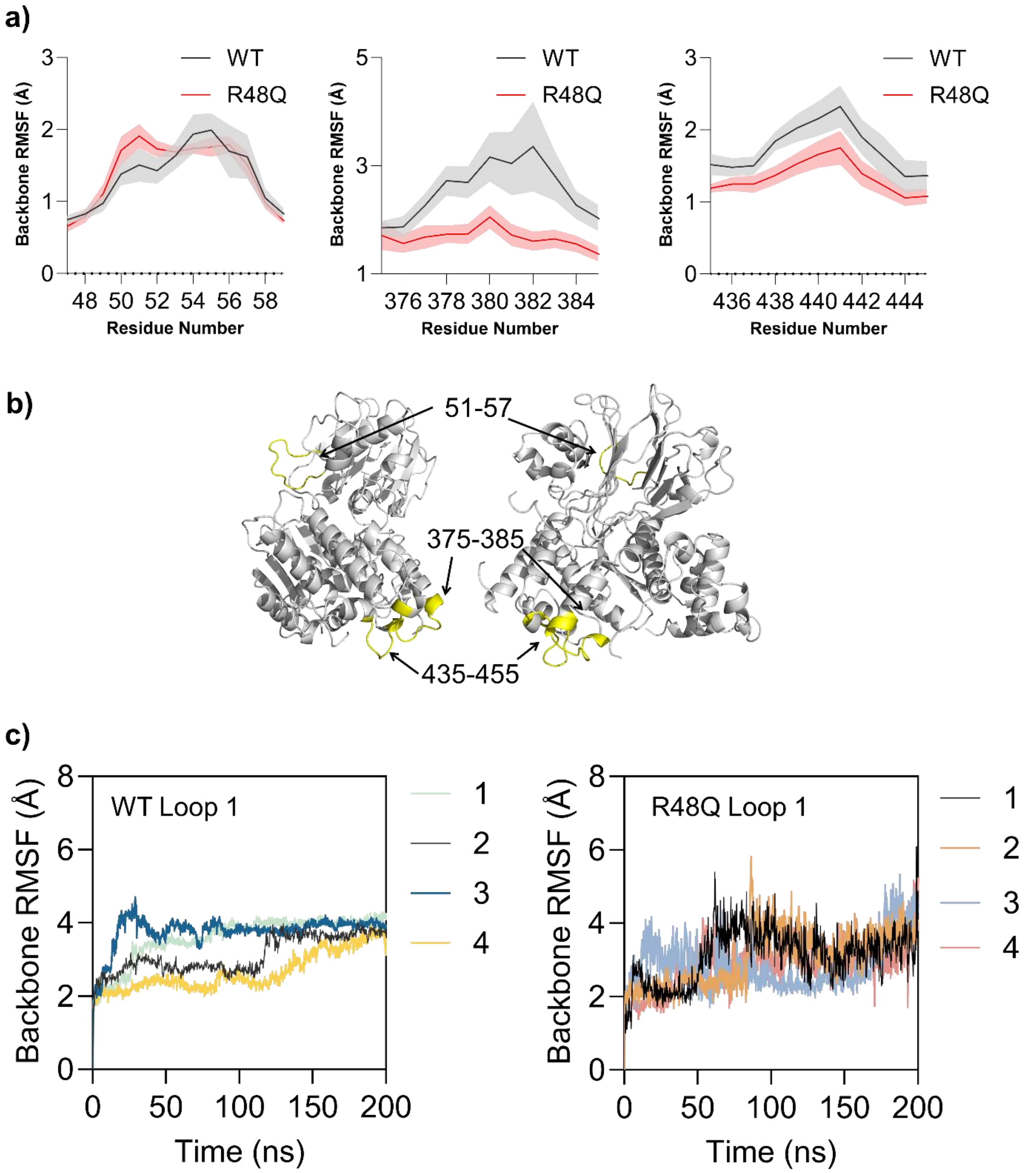


**Fig. S19. The RMSF comparison between WT and R48Q.** a) Three regions exhibiting different dynamical motion in the R48Q variant are observed near Loop 1 (48-58) and the synthetase domain (375-385), and the C-terminal tail (435-445). b) structural presentation of the highlighted region in the RMSF plot of a. c) Backbone RMSD plot for the four replicates in the MD trajectories at Loop 1. Plots show the time-dependent RMSD fluctuation (Å) of Loop 1 in WT (left) and the R48Q variant (right) over four 200-ns MD trajectories. Each simulation was performed in four replicates, indicated in different colors.

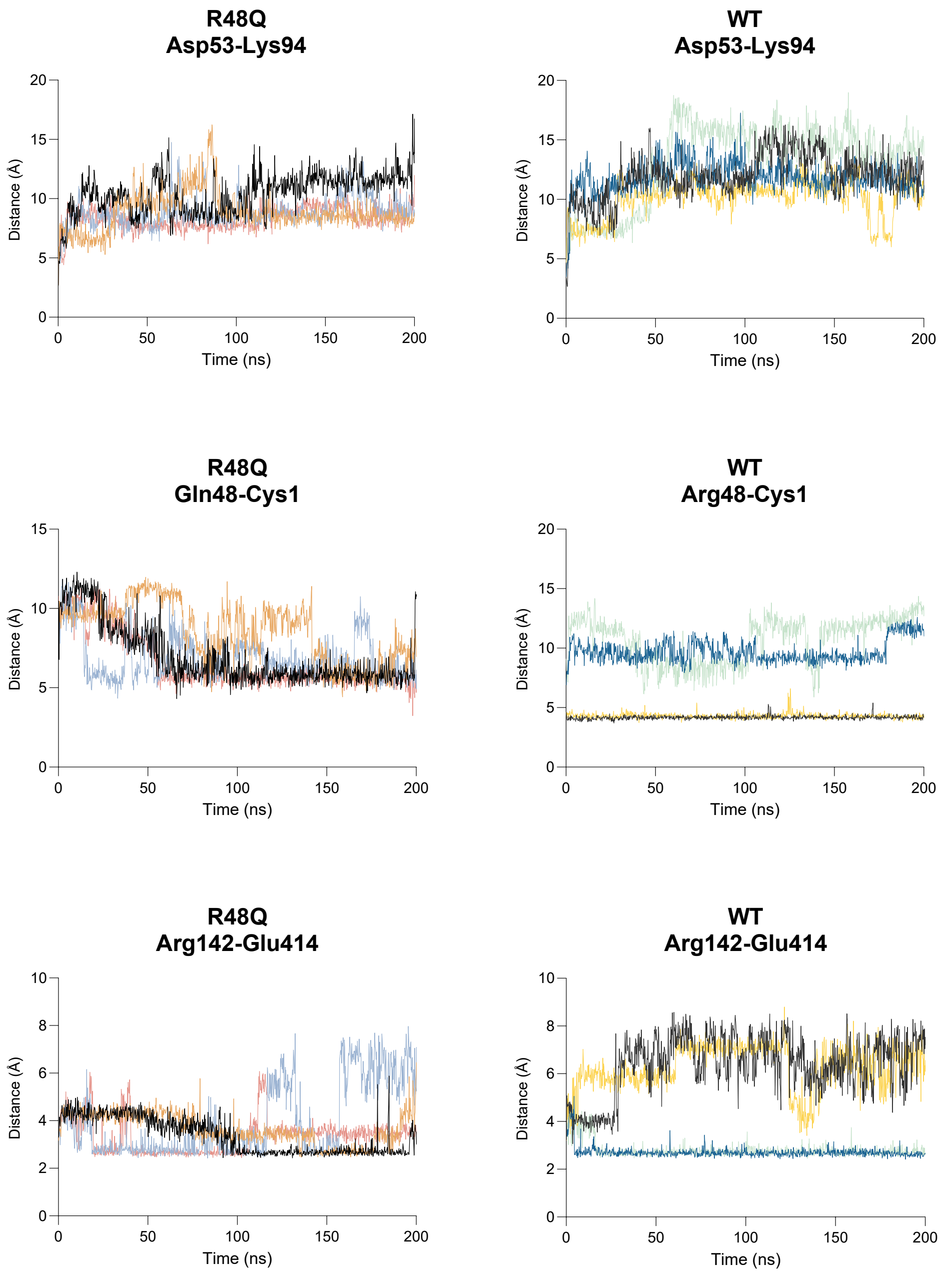


**Fig. S20. Analyses of distances between residues from MD simulations of WT and the R48Q variant.** Plots show the time-dependent distances (Å) between selected residue pairs over four 200-ns MD trajectories for the R48Q variant (left) and WT (right). Top, distances between Asp53-Lys94 report on Loop 1 dynamics in the N-terminal domain. Middle, distances between Gln48/Arg48 and Cys1 reflect interactions near the N-terminal catalytic site. Bottom, distances between Arg142-Glu414 probe long-range coupling between the ammonia tunnel gate (Arg142) and the C-terminal synthetase domain. Together, these results illustrate that R48Q alters local Loop 1 interactions and propagates dynamic changes to distal functional regions.

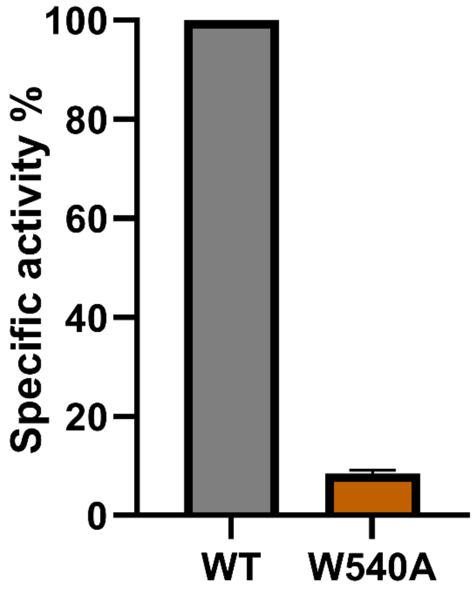


**Fig. S21. W540A exhibits impaired catalytic activity in producing pyrophosphate.** Pyrophosphate production is greatly decreased in the W540A variant, which shows that the interaction between W540 and the hydrophobic pocket in the C-terminal domain is crucial for the catalytic activity at the C-terminal active site.

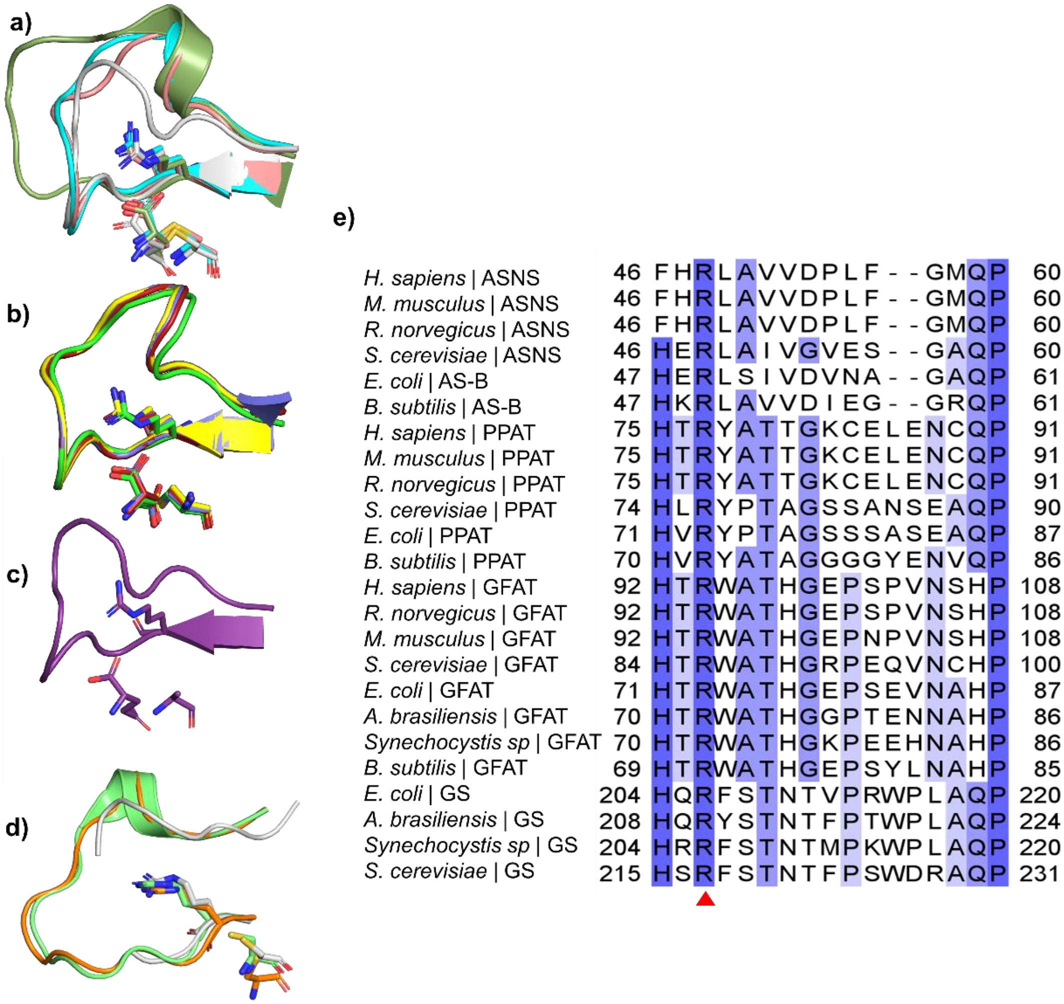


**Fig S22. Structural superimposition and sequence alignment of Loop 1 from apo and ligand-bound structures of Class II GATase family members.** a) Superimposition of Loop 1 of DON modified structures of human asparagine synthetase (ASNS, gray, PDB: 6GQ3), *E. coli* amidophosphoribosyltransferase (PPAT, cyan, PDB: 1ECG), *E. coli* glutamine-fructose-6-phosphate aminotransferase (GFAT, salmon, PDB: 2J6H), and *Synechocystis sp*. glutamate synthase (GS, smudge, PDB: 1OFE). b) Superimposition of Loop 1 of glutamate-bound structures of *E. coli* glutamine-fructose-6-phosphate aminotransferase (GFAT, slate, PDB: 1XFF), human glutamine-fructose-6-phosphate aminotransferase (GFAT-1 in complex with C-terminal ligand, glucosamine-6-Phosphate, yellow, PDB: 6SVO), human glutamine-fructose-6-phosphate aminotransferase (GFAT-1 in complex with glucose-6-phosphate, green, PDB: 6R4E) and human glutamine-fructose-6-phosphate aminotransferase (GFAT-1 in complex with glucose-6-Phosphate and UDP-GalNAc, raspberry, PDB: 6SVM) c) Superimposition of Loop 1 of the substrate-bound structure of *E. coli* glutamine-hydrolyzing asparagine synthetase (AsnB, purple, PDB: 1CT9). d) Superimposition of Loop 1 of the apo human asparagine synthetase (ASNS, gray, PDB: 8SUE) and substrate/product free holoenzyme structure of *Synechocystis* sp. glutamate synthase (GS, lime green, PDB: 1LM1) and *Azospirillum brasilense* glutamate synthase (GS, orange, PDB: 6S6S). e) Loop 1 is conserved across Class II GATases, including ASNS, PPAT, GFAT, and GS. Conserved residues are highlighted in blue, with the intensity of the color corresponding to the degree of conservation for Class II GATase members. The conserved arginine residue across all enzymes in several organisms is marked with a red triangle.

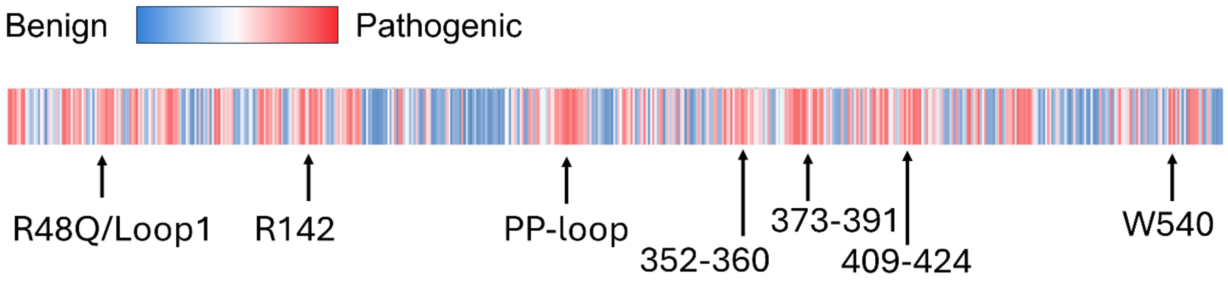


**Fig. S23. AlphaMissense pathogenicity predictions across the ASNS protein sequence.** Color-coded heatmap representation of AlphaMissense predictions for all possible missense variants in human ASNS. Variants predicted to be pathogenic are shown in shades of red, whereas benign predictions are shown in blue. These predictions align with the residue network identified in this study, in which many of the sites predicted to be functionally critical overlap with conserved structural motifs and interdomain communication residues. Pathogenic predictions cluster in regions involved in the N-to-C-terminal domain interface and the catalytic cores, supporting the hypothesis that disruption of dynamic residue networks contributes to ASNSD.

**Table S1. Clinically identified ASNSD-associated variants.** All residue numbers were shifted by one due to the post-translational removal of the initiator methionine. The reference sequence is NM_001673.5.

| ASNS Variants | Molecular Consequence | Protein Change * | Zygosity | Clinical Notes | Ref |
|---|---|---|---|---|---|
| c.17C>A & c.1648C>T | Missenses | A5E & R549C | Compound heterozygous | Severe developmental delay, progressive microcephaly, partial complex seizures, axial and appendicular hypotonia, decreased cerebral volume and size of pons, simplified gyri. | (16) |
| c.97C>T & c.1031-2_1033del | Missense & splicing site | R32C | Compound heterozygous | Severe brain dysplasia | (17) |
| c.144C>A | Missense | H47Q | Homozygous | Developmental delay, epilepsy, spasticity | (18) |
| c.146G>A | Missense | R48Q | Homozygous | Developmental delay, epilepsy, microcephaly, brain atrophy | (19) |
| c.224A>T | Missense | N74I | Homozygous | Developmental delay, epilepsy, microcephaly, brain atrophy | (20) |
| c.224A>G & c.413A>C | Missenses | N74S & D137A | Compound heterozygous | Spasticity, hyperreflexia, microcephaly, decreased size of pons | (21) |
| c.224A>G & c.1612A>G | Missenses | N74S & M537V | Compound Heterozygous | Development delay, epilepsy, microcephaly, brain atrophy | (22) |
| c.368T>C & c.1649G>A | Missenses | F122S & R549H | Compound Heterozygous | Developmental delay, epilepsy, brain atrophy, decreased size of pons, gyral simplification, hyperreflexia, microcephaly, visual impairment, axial hypotonia | (23) |
| c.434T>C & c.740T>G | Missenses | L144S & L246W | Compound heterozygous | Epilepsy, hyperreflexia, axial hypotonia, developmental delay | (24) |
| c.478del & c.1283A>G | Frame shift & Missense | E159KfsTer8 & Y427C | Compound heterozygous | Profound microcephaly, lissencephaly, epilepsy, pachygyria, cerebral and cerebellar volume loss, feeding difficulties, muscle contractures, profound cognitive impairment, cortical blindness, and dysmorphic features | (25) |
| c.538T>A | Missense | F179I | Homozygous | Progressive microcephaly, Axial hypotonia, seizures | (26) |
| c.601delA & c.1165G>C | Frame shift & Missense | M200fs & E388Q | Compound heterozygous | Developmental delay, microcephaly, cortical blindness, seizure | (27) |
| c.614A>C & c.1192dupT | Missense & frame shift | H204P & Y397LfsTer4 | Compound heterozygous | Developmental delay, epilepsy, microcephaly, brain atrophy | (28) |

| | | | | | |
|---|---|---|---|---|---|
| c.728T>C & c.1097G>A | Missenses | V242A & G365E | Compound heterozygous | Developmental delay, microcephaly, cortical blindness, seizure | (29) |
| c.788C>T | Missense | S262F | Homozygous | Developmental delay, epilepsy and microcephaly | (30) |
| c.866G>C & c.1010C>T | Missenses | G288A & T336I | Compound heterozygous | Severe developmental delays, epilepsy, brain atrophy | (31) |
| c.1019G>A | Missense | R339H | Homozygous | Microcephaly, cerebral atrophy | (32) |
| c.1084T>G | Missense | F361V | Homozygous | Seizures, Hypsarrhythmia, microcephaly | (16) |
| c.1108C>T | Missense | L369F | Homozygous | Developmental delay, epilepsy, microcephaly, brain atrophy | (33) |
| c.1118G>T & c.1556G>A | Missense | G372V & R518H | Compound heterozygous | Developmental delay, epilepsy, microcephaly, brain atrophy | (34) |
| c.1160 A>G | Missense | Y375C | Homozygous | Progressive microcephaly, Axial hypotonia, seizures | (35) |
| c.1138G>T | Missense | A379S | Homozygous | Developmental delay, epilepsy, microcephaly | (36) |
| c.1193A>G | Missense | Y397C | Homozygous | Developmental delay, Hyptertonia, Spastic quadriplegia, seizure, severe microcephaly thin corpus callosum, brain atrophy, decreased size of pons, simplified gyral pattern | (37) |
| c.1211G>A | Missense | R403H | Homozygous | Congenital microcephaly, progressive postnatal microcephaly, spasticity, seizures | (21) |
| c.1219C>T | Nonsense | R406Ter | Homozygous | Epilepsy, hyperreflexia | (38) |
| c.1424C>A | Missense | T474N | Homozygous | Developmental delay, epilepsy, microcephaly, gyral simplification | (20) |
| c.1424C>T<br><br>c.666_ 667delCT | Missense & frame shift | T474I<br><br>L222LfsTer5 | Compound Heterozygous | Epilepsy, developmental delay, congenital visual impairment | (39) |
| c.1439C>T& c.1648C>T | Missenses | S479F & R549C | Compound heterozygous | Developmental delay, epilepsy, microcephaly, gyral simplification | (40) |
| c.1466T>A & c.1623_1624del | Missense & frame shift | V488D & W540CfsTer5 | Compound heterozygous | Epilepsy, hyperreflexia, axial hypotonia, developmental delay | (24) |
| c.1648C>T | Missense | R549C | Homozygous | Progressive microcephaly, developmental delay, Axial hypotonia | (16) |
| c.1649G>A | Missense | R549H | Homozygous | Developmental delay, epilepsy, spasticity, visual impairment, decreased size of pons | (21) |
| c.1648C>T | Missense | R549C | Homozygous | Developmental delay, epilepsy, microcephaly, brain atrophy | (16) |
| c.1476+1G>A | Splice donor | n/a | Homozygous | Developmental delay and cerebellar hypoplasia | (41) |

**Table S2. Cryo-EM data collection, refinement, and validation statistics**

| | | |
|---|---|---|
| **Data collection** | | |
| | ASNS (WT) | ASNS-ASX173 (R48Q) |
| Magnification | ×105,000 | ×105,000 |
| Voltage (kV) | 300 | 300 |
| Electron exposure (e/Å$^2$) | 59.45 | 59.45 |
| Defocus range (μm) | -0.8 to -1.8 | -0.8 to -1.8 |
| Pixel size (Å) | 0.411 | 0.411 |
| **Data processing** | | |
| | **EMD-75273, PDB 10LS** | **EMD-75275, PDB 10LT** |
| Symmetry imposed | C2 | C2 |
| Initial particle images (no.) | 3,027,978 | 4,341,491 |
| Final particle images (no.) | 90,670 | 76,456 |
| Map resolution (Å) | 2.78 | 2.81 |
| FSC threshold | 0.143 | 0.143 |
| Map resolution range | 2.4-3.2 | 2.4-3.2 |
| Map sharpening B factor (Å$^2$) | 93.1 | 95.6 |
| **Refinement** | | |
| Model resolution (Å$^2$) | 2.0 | 2.0 |
| Model composition | | |
| Non-hydrogen atoms | 8466 | 8414 |
| Protein residues | 1043 | 1037 |
| Ligands. | 0 | 0 |
| B-factors (min/max/mean) | | |
| Protein | 16.53/115.46/45.42 | 23.28/99.04/44.96 |
| Ligand | | |
| r.m.s.d. deviations | | |
| Bond length (Å) | 0.003 | 0.003 |
| Bond angles (°) | 0.594 | 0.594 |
| Validation | | |
| MolProbility score | 2.5 | 2.54 |
| Clashscore | 11.82 | 11.43 |
| Rotamer outliers (%) | 5.15 | 5.52 |
| CaBLAM outliers (%) | 2.85 | 3.26 |
| Ramachandran plot (%) | | |
| Outliers | 0 | 0 |
| Allowed | 5.43 | 5.95 |
| Favored | 94.57 | 94.05 |
